# The E-cadherin-ESR1-GRPR axis defines a sex-specific metastatic pathway in melanoma

**DOI:** 10.1101/2022.12.02.518844

**Authors:** Jérémy H. Raymond, Marie Pouteaux, Valérie Petit, Zackie Aktary, Flavie Luciani, Maria Wehbe, Patrick Gizzi, Claire Bourban, Igor Martianov, Irwin Davidson, Catherine-Laure Tomasetto, Florence-Mahuteau Betzer, Béatrice Vergier, Lionel Larue, Véronique Delmas

## Abstract

Although tremendous progress has been made in understanding the mechanisms leading to cancer, those governing metastases are still poorly understood. E-cadherin (Ecad) is a cell-cell adhesion molecule essential for tissue homeostasis, and its loss often correlates with the dissemination of human cancers. However, whether and how the loss of Ecad triggers the full metastatic program is largely unknown. Here, we show that the loss of Ecad promotes melanoma lung metastases in females. The loss of Ecad, after the induction of estrogen receptor α (ERα) expression, activates gastrin-releasing peptide receptor (GRPR) expression. GRPR promotes cellular processes essential for metastasis formation through G□_q_ and YAP1 signaling and its pharmacological inhibition reduces metastasis *in vivo*. This study reveals an Ecad-ERα-GRPR metastatic sex dimorphism axis in melanoma that is conserved in human breast cancer and provides proof of concept that the G-coupled receptor GRPR is a therapeutic target for metastasis.

## Introduction

Recent therapeutic advances have made it possible to combat metastases, but several major problems remain. In addition to non-responsive patients or innate/acquired resistance to therapies, the efficacy of treatments for metastases is still limited. The metastatic process is complex, requiring multiple steps, such as the escape of cells from the primary tumor, intravasation and survival in the bloodstream or lymphatic circulation, extravasation and invasion of the metastatic site, proliferation, and adaptation of the cells to new environments that are not always favorable. The identification of unique drivers of the metastatic processes capable of orchestrating all the steps leading to the colonization of distant organs by tumor cells is still limited, probably due to the intrinsic complexity of said processes and the fact that most mechanistic studies have been performed *in vitro* or in xenograft models. Cutaneous melanoma is a very aggressive cancer with high metastatic potential that results in the death of half of all patients, despite novel therapies ^1^. The incidence of metastatic melanoma continues to increase with each decade and new therapeutic options must be developed to combat the progression of the disease. Melanoma is highly heterogeneous among patients and within the tumor itself, which may explain treatment failures. Exploration of underlying characteristics among patients grouped by age, ethnicity, or sex would likely provide additional clues for new therapeutic options. Melanoma develops through the acquisition of genetic and epigenetic changes of oncogenes and tumor suppressors, not to mention its particular plasticity including the phenotypic switch from the proliferative to invasive cell state or cell differentiation ^2-6^.

Epithelial-mesenchymal transition (EMT) is a mechanism of cellular plasticity that has been reported in carcinoma, as well as in non-epithelial tumors, including melanoma, following a pseudo-EMT. EMT is a highly dynamic process and is promoted by multiple oncogenic signaling pathways and a variety of signals from the tumor microenvironment, such as growth factors and hormones ^7, 8^. EMT is associated with the acquisition of invasive and metastatic properties, as well as stemness. Loss of the cell-cell adhesion molecule, E-cadherin (Ecad), is considered to be the hallmark of EMT, resulting in the loss of cell-cell adhesion and cell-matrix interactions during normal and pathological development. By regulating a wide range of cellular functions, including gene expression, Ecad is a key protein in the establishment of epithelial architecture ^9–11^. In the skin, Ecad regulates melanocyte homeostasis and is the primary adhesion molecule that mediates melanocyte-keratinocyte interactions. Its loss is associated with melanocyte detachment from the basal to suprabasal layer of the epidermis and the pathogenesis of vitiligo ^12, 13^. In carcinomas, *CDH1*, encoding Ecad, is considered to be a tumor suppressor gene and its loss promotes carcinoma progression ^14, 15^. The role of Ecad during melanomagenesis is still unclear, as only correlative studies have been performed. Its loss is observed in 40% of primary melanoma and correlates with the presence of metastasis ^16, 17^. Surprisingly, how the loss of Ecad affects the signaling network and cellular properties in melanoma has never been truly addressed, despite its importance in other cancers.

Here, we show that the loss of Ecad in melanocytes does not affect tumor initiation but rather promotes tumor progression in a spontaneous mouse model, specifically in females. Using transcriptomics, we identify a marked increase in the level of the gastrin-releasing peptide receptor (*Grpr*) in female melanoma lacking Ecad. Upon the binding of its ligand, GRPR acts as a driver of the metastatic process to confer melanoma cells with the capacity to resist anoïkis and to promote invasion, growth, and lung colonization, all of which are abolished in the presence of a GRPR antagonist. Finally, we discovered a signaling mechanism by which the loss of Ecad induces transcription of the estrogen receptor ERα, which, in turn, regulates *GRPR* expression in melanoma, as well as breast carcinoma. These data highlight a previously unidentified sex-specific Ecad-ERα-GRPR metastatic axis that opens a new avenue for targeted therapy for personalized medicine.

## Results

### The loss of Ecad increases metastatic potential without affecting the initiation of melanoma

We established a mouse model with a conditional deletion of *Cdh1* in melanocytes (*Tyr*::*CreA*/°; *Cdh1*^F/F^) to decipher the role of Ecad in melanomagenesis. Even after two years, no melanoma had developed, showing that the loss of *Cdh1* alone is insufficient to induce melanoma formation. To initiate melanoma, we associated the loss of *Cdh1* with the *NRAS^Q61K^* mutation in melanocytes using the well-characterized *Tyr::NRAS^Q61K^*/°; *Cdkn2a*^+/-^ melanoma model ^18^. The choice of using an NRAS over BRAF mutation was based on the following observations in humans: the repression of *CDH1* is more frequently associated with NRAS than BRAF mutations and no targeted therapy for an NRAS-mutated melanoma is yet available (TCGA database Figure S1A). *Tyr::NRAS^Q61K^/*°; *Cdkn2a*^+/-^; °/°; *Cdh1*^F/F^ (called Ecad from hereon) and *Tyr::NRAS^Q61K^*/°; *Cdkn2a*^+/-^; *Tyr::CreA*/°; *Cdh1*^F/F^ (called ΔEcad from hereon) mice were followed for melanoma development. The *Cdh1* loss did not modify the kinetics of appearance of the primary melanoma (Figure 1A). The median time to onset and the frequency of apparition were similar between Ecad and ΔEcad mice (Figure S1B and S1C). There were no differences between the two groups of mice in terms of Ecad status or sex (Figure S1D-S1G). However, the incidence of lung melanoma metastasis was significantly higher in the absence of *Cdh1*: 18% in the colonized lungs of Ecad versus 49% in those of ΔEcad mice (Figure 1B-1D). The number of metastases per lung was also seven times higher in the ΔEcad group compared to the Ecad group (Figure 1E). Surprisingly, analysis of metastasis occurrence by sex showed the frequency and number of metastases to be much higher in ΔEcad females than ΔEcad males (Figure 1F and 1G). The number of both micro- and macro-metastasis was higher in ΔEcad females than ΔEcad males (Figure 1H and 1I). No sex-based differences were observed in Ecad mice. Thus, the loss of Ecad influences the metastatic potential of melanoma in a sex-dependent manner.

**Figure 1.**
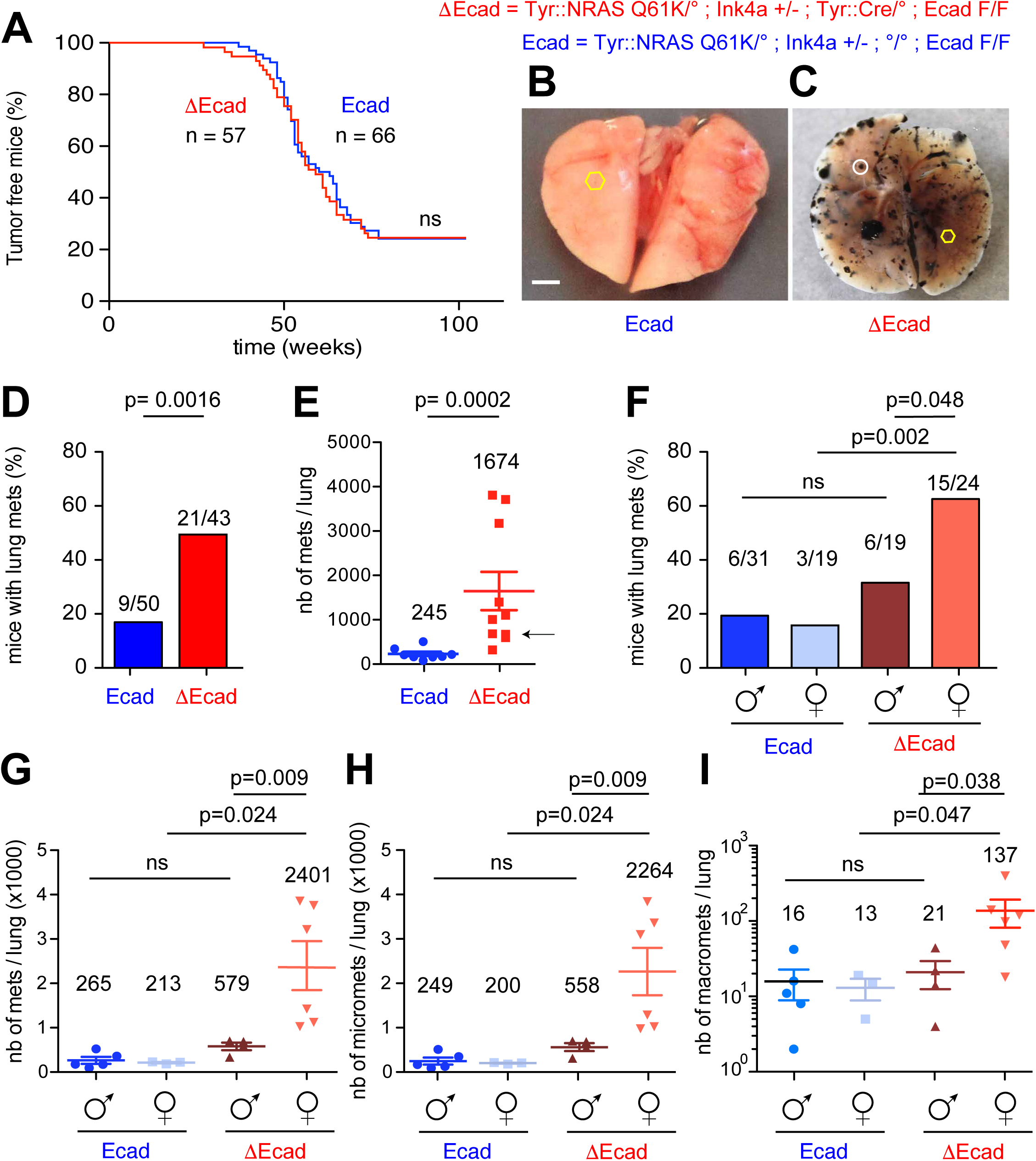
Loss of Ecad in primary melanoma promotes lung melanoma metastasis in females. (**A**) Kaplan-Meier curves of melanoma-free Ecad mice (n = 66) and ΔEcad mice (n = 57). The curves were not significantly different according to the log-rank method. Representative images of the lungs of Ecad (**B**) and ΔEcad (**C**) mice with primary melanoma are depicted. Yellow hexagons and white circles show micro-metastases and macro-metastases, respectively. Scale bar= 2 mm. (**D, F**) Frequency of mice with lung metastasis and (**E, G**) mean number of lung melanoma metastases in mice according to Ecad status (**D, E**) and Ecad and sex status (**F, G**). Micro-(**H**) and macro-(**I**) metastasis quantifications according to Ecad and sex status. Micro- and macro-metastases are defined as metastases visible under the binocular microscope and naked eye, respectively.

### The loss of E-cadherin upregulates GRPR expression

To understand the cellular and molecular mechanisms induced by the loss of Ecad, we performed RNA-seq analysis comparing eight primary tumors per genotype and sex. Grpr expression was markedly upregulated in ΔEcad female melanoma relative to that of all other genotypes, including ΔEcad males (Figure 2A, Tables S1-S4). The relevance of Grpr in conferring aggressive behavior to tumor cells was supported by the high correlation (Pearson r = 0.47) of Grpr expression with the invasive score in human melanoma (Figure 2B). These results were confirmed by quantitative RT-PCR analysis, showing Grpr to be highly expressed in only ΔEcad female tumors (Figure 2C). Grpr overexpression in female melanomas was independent of the Tyr::Cre line used, as we obtained the same results with the Tyr::CreB line ^19^ crossed with *Tyr::NRAS^Q61K^*; *Cdkn2a*^+/-^; *Cdh1*^F/F^ mice. In addition, Tyr::CreA expression did not induce Grpr expression in other genetically engineered melanomas tumors derived from, for example, *Tyr::NRAS^Q61K^*; *Tyr::CreA*; *Pten^F/F^*mice (Figure 2D).

**Figure 2.**
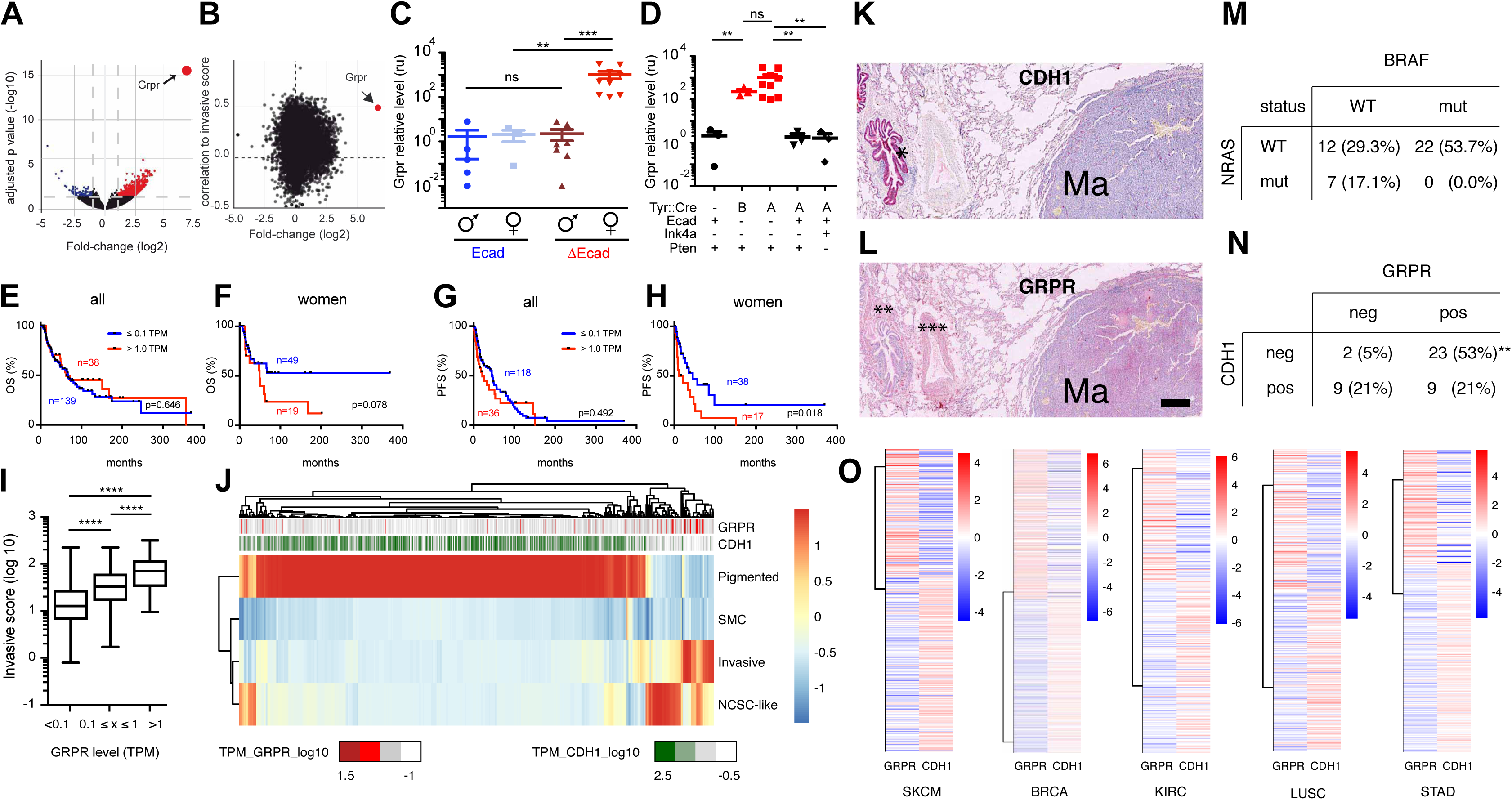
Loss of Cdh1 induces Grpr expression. (**A**) Volcano plot depicting the differential expression between female Ecad and female ΔEcad tumors with Grpr indicated in red. (**B**) Plot of the differential gene expression (fold change, x-axis) with the Pearson correlation to “invasive” score (y-axis). The fold change in x-axis was calculated by the differential gene expression between female ΔEcad and female Ecad tumors. The “invasive” score was calculated by averaging the expression of gene markers of the “invasive” state (CYR61, WNT5A, TNC, TCF4, LOXL2, and AXL) for each TCGA-SKCM tumor. The correlation of each genes expressed in the TCGA-SKCM tumors to the “invasive” score in y-axis was calculated using Pearson test. mRNA levels of Grpr as measured by quantitative reverse transcriptase-PCR (q-RT-PCR) in (**C**) Ecad and ΔEcad melanoma in males and females and in (**D**) different Tyr::NRAS^Q61K^ transgenic melanoma genetic backgrounds. Tyr::CreA represented by the “A” and Tyr::CreB by the “B” mouse lines ^19^, Ecad^+/+^ by “+”, Ecad^F/F^ by “-“, Ink4a^+/-^ by “-“ and Ink4a^+/+^ by “+”, Pten^+/+^ by “+”, and Pten^F/F^ by “-“. Kaplan-Meier curves of the overall survival (OS) of (**E**) all patients and of (**F**) women only according to GRPR expression (low/absent ≤ 0.1 TPM and expressed > 1 TPM). Kaplan-Meier curves of the progression-free survival (PFS) of (**G**) all patients and of (**H**) female patients according to GRPR expression (low/absent ≤ 0.1 TPM and expressed > 1 TPM). (**I**) Invasive score according to GRPR level in melanomas from the TCGA-SKCM database. Spearman coefficient r = 0.35. (**J**) Heatmap clustering TCGA-SKCM samples according to the main expressed cell-state signatures: “Pigmented”, “SMC” (starved-like melanoma cells), “Invasive”, and “NCSC-like” (neuronal crest cell-like). GRPR and CDH1 levels are indicated as a color gradient from dark colors (higher expression, red for GRPR and green for CDH1) to white (lower expression). Immunohistochemistry staining for (**K**) CDH1 and (**L**) GRPR in human lung melanoma metastases. *Bronchi as an internal positive control for ECAD staining and ** and ***smooth muscle as an internal positive control for GRPR. Metastasis classification according to (**N**) NRAS and BRAF status and (**M**) CDH1 and GRPR expression of the lung metastasis samples. (**O**) Heatmap of GRPR and CDH1 expression in melanomas and different carcinomas extracted from the TCGA database. SKCM: skin cutaneous melanoma, BRCA: breast invasive carcinoma, KIRC: kidney renal clear cell carcinoma, LUSC: lung squamous cell carcinoma, and STAD: stomach adenocarcinoma.

Before addressing the role of GRPR, we sought to validate findings from our mouse model in human melanoma. We first analyzed the TCGA database and compared the overall survival (OS) and progression-free survival (PFS) of patients with a melanoma expressing *GRPR* (> 1TPM) relative to melanomas not expressing *GRPR* (≤ 0.1TPM). While the expression of *GRPR* appeared to influence the OS of women (p = 0.076), it had a truly dramatic impact on PFS (p = 0.018), confirming a role for GRPR in disease progression in women (Figure 2F and 2H). However, this difference was no longer observable when men were included (Figure 2E and 2G). Moreover, the analysis of tumor transcriptomic data showed that *GRPR* expression correlated with an “invasive” phenotype (r = 0.47) (Figure 2I and 2J) and that its expression was highly enriched in tumors with low *CDH1* expression (p = 0.002) (Figure 2J). Aside from the strong correlation with the “invasive” phenotype, *GRPR*-expressing melanomas clustered with the “neuronal crest stem cell (NCSC) like” phenotype (r = 0.32) and inversely with the pigmentation phenotype (r = -0.31) (Figure 2J and S2A-S2C). In addition, we evaluated GRPR expression by immunohistochemistry in lung melanoma metastasis derived from patients harboring a number of different driver mutations (Figure 2K-2M); 74% of the lung metastases expressed GRPR, with the most frequent profile being Ecad-neg/GRPR-pos (Figure 2N). These results indicate that the mouse model we developed is relevant and reflects human melanoma pathology. Of note, we observed the loss of Ecad expression in 58% of lung melanoma metastases, indicating that expression (or re-expression) of Ecad in metastatic sites, as is often suggested, is not necessarily required for melanoma cell colonization in the lung.

The anticorrelation between *CDH1* and *GRPR* expression observed in melanoma (SKCM) was also observable in carcinomas, such as breast cancer (BRCA), kidney cancer (KIRC), lung cancer (LUSC), and stomach cancer (STAD), according to the TCGA database. This indicates that the regulation of GRPR expression by ECAD could be common to many cell types and/or cancers (Figure 2O).

As the endogenous ligand of GRPR, GRP, is particularly expressed in the lungs of rodents and humans (Figure S2D), we hypothesize that the activation of GRPR by GRP in the lungs could activate cellular mechanisms, such as invasion and cell growth, to promote the formation and establishment of metastases.

### GRPR expression promotes cellular mechanisms essential for melanoma progression

We next wanted to assess whether the mechanism(s) that lead to lung colonization by ΔEcad melanoma cells is cell-autonomous and/or mainly due to cell-cell adhesion defects related to Ecad loss. Thus, we dissociated primary tumors and injected the cells into the tail veins of C57BL6/J mice. ΔEcad melanoma cells were able to colonize the lungs within 30 days, in contrast to Ecad melanoma cells (Figure S3A). Thus, ΔEcad melanoma cells (which express GRPR) were able to efficiently colonize the lungs and effectively adapted to the lung environment. We next engrafted ΔEcad and Ecad female tumors into female C57BL/6J mice. The ΔEcad female tumors grew significantly more efficiently than the Ecad female tumors (Figure S3B). Thus, efficient lung colonization and rapid growth after grafting support the cell-autonomous aggressiveness of ΔEcad female melanomas. As we hypothesized that the interaction of GRP with GRPR is central to melanoma progression, we next evaluated the effect of GRP stimulation on GRPR for the fundamental cellular processes required for the metastatic pathway.

Ecad and ΔEcad female and male melanomas were established as cell lines in culture from the mouse primary melanoma; consistent with the mouse primary melanomas, only ΔEcad female melanoma cell lines produced GRPR (Table S5). GRP stimulation induced the cell growth of mouse (1057) and human (MDA-MB-435S) melanoma cell lines expressing GRPR (GRPR^pos^) (Figure 3A and 3B). Co-treatment of these cells with the agonist GRP and antagonist RC-3095 (RC) blocked GRP-induced growth. Note that RC is a partial agonist that acts as an antagonist in the presence of an agonist (GRP) and as a weak agonist without GRP, explaining the slight growth induction of cells treated with RC alone ^20^. FCS was used as a positive control of induction. The regulation of cellular growth by GRP was confirmed using another cell line expressing GRPR, 1064 (Figure S3D). As expected, GRP treatment had no effect on GRPR-negative (GRPR^neg^) mouse (1181 and 1014) and human (501mel) melanoma cell lines (Figure 3C, 3D, and S3E). We addressed the specificity of GRPR induction by GRP by transfecting GRPR into GRPR^neg^ 1181, 1014, and 501mel melanoma cells. Both murine and human melanoma cells transfected with Grpr responded to GRP induction and RC inhibition (Figure 3E, 3F, and S3F).

**Figure 3.**
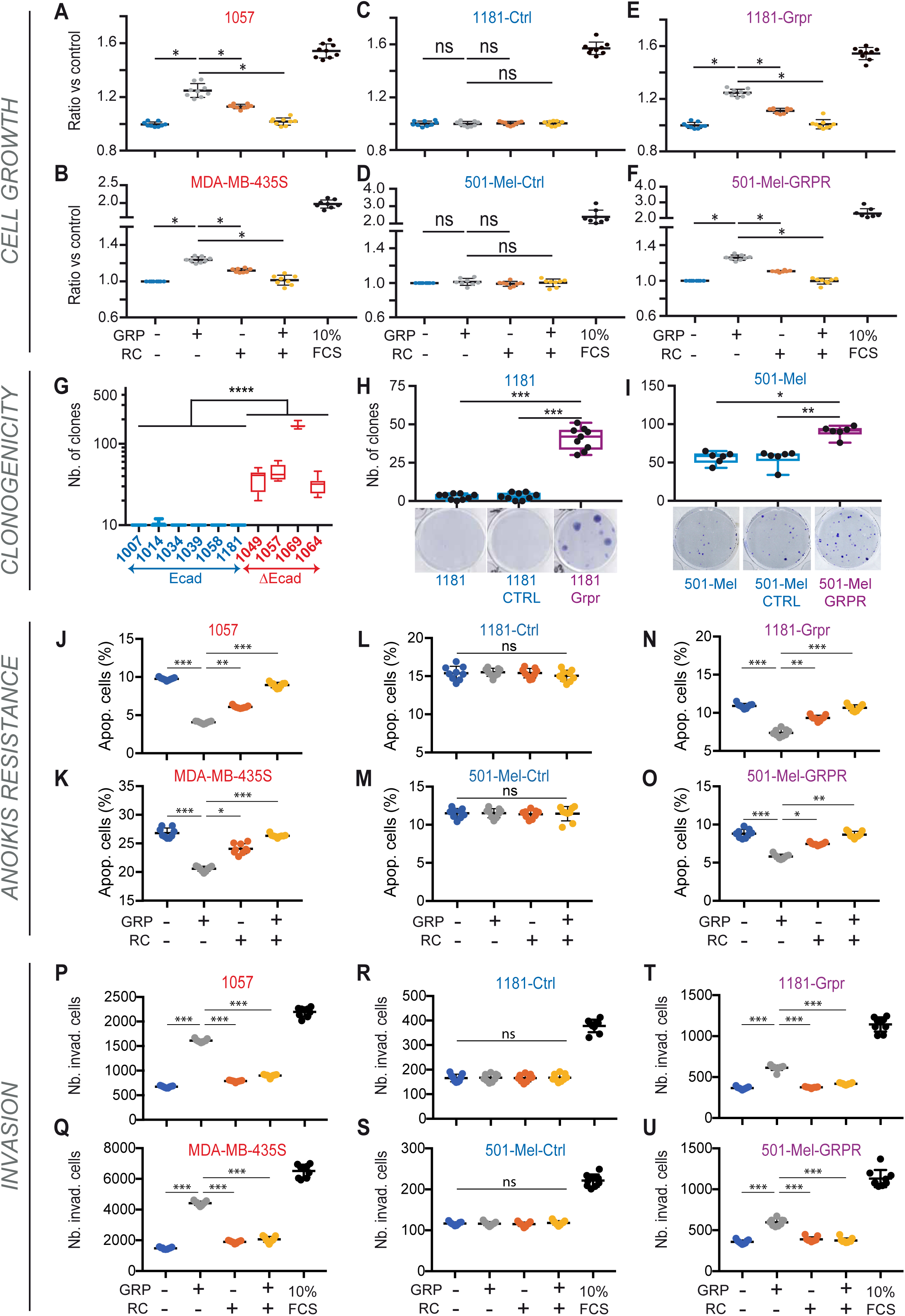
GRPR promotes growth, survival, and invasiveness of melanoma cells. (**A-F**) GRPR activation promotes cell growth in vitro. Mouse (**A**) 1057, (**C**) 1181-Ctrl, and (**E**) 1181-Grpr and human (**B**) MDA-MB-435S, (**D**) 501-Mel-Ctrl, and (**F**) 501-Mel-GRPR were used. 1057 and MDA-MB-435S are GRPR^pos^ cells and 1181 and 501-Mel are GRPR^neg^ cells. (**G-I**) GRPR expression induces cell clonogenicity. (**G**) Colony formation 10 days after seeding 500 melanoma cells. In blue, Ecad^pos^/GRPR^neg^ cells (Ecad) and in red, Ecad^neg^/GRPR^pos^ cells (ΔEcad). (**H-I**) Clonogenic growth of the (**H**) 1181 mouse and (**I**) 501-Mel human GRPR^neg^ melanoma cell line: left, parental cells; middle, control cell line; right, exogenous GRPR expression. Data are presented as the mean ± SD (**J-O**) GRPR activation induces resistance to anoikis. Percentage of apoptotic (Apop.) cells after 48 h of growth without attachment to the matrix of (**J**) 1057, (**K**) MDA-MB-435S, (**L**) 1181-Ctrl (**M**) 501-Mel-Ctrl, (**N**) 1181-Grpr, and (**O**) 501-Mel-GRPR cells. Note that the number of apoptotic cells is reduced when GRPR is expressed in 1181 (15.4% *vs*. 10.9%, p = 0.0001) and 501-Mel (11.5% *vs*. 8.1%, p = 0.0017) cells. (**P-U**) GRPR activation induces Matrigel invasion. GRPR^pos^ cells (**P**) 1057, (**Q**) MDA-MB-435S, (**T**) 1181-Grpr, (**U**) 501-Mel-GRPR and GRPR^neg^ (**R**) 1181-Ctrl, (**S**) 501-Mel-Ctrl. Number (Nb) of invading cells (invad.) (**A-F, J-U**) Twenty-four hours after seeding, cells were incubated in low serum for 18 h in 0.5% FCS for murine (**A, C, E, J, L, N, P, R, T**) and 1% for human (**B, D, F, K, M, O, Q, S, U**) cell lines. Then, cells were treated with vehicle (lane 1), 10 nM GRP (lane 2), 1 µM RC (lane 3), 10 nM GRP + 1 µM RC (lane 4), or 10% FCS (lane 5, when present). The read out was performed 24 h later for invasion, and 48 h later for growth and resistance to anoikis. At least three independent biological experiments were performed for each assay. Statistical analysis was performed by ANOVA. Data are presented as the mean ± sd. ns = not significant, *p < 0.05, **p < 0.01, ***p < 0.001, and ****p < 0.0001.

GRPR^pos^ (1049, 1057, 1069, and 1064) mouse melanoma cells were able to grow efficiently as single cells in culture, whereas GRPR^neg^ (1007, 1014, 1034, 1039, 1058, and 1181) cells did not (Figure 3G). Ectopic expression of GRPR in mouse (1181-Grpr and 1014-Grpr) or human (501-Mel-GRPR) GRPR^neg^ melanoma cells significantly promoted clonogenic growth (Figure 3H, 3I, and S3C).

We performed RNA-seq to assess the differentially expressed genes induced by GRPR activation. The analysis showed that GRPR activation by GRP is likely to lead to the resistance of anoikis (Figure S3G-S3I), as well as the promotion of invasion (Figure S3J-S3L) in murine and human GRPR^pos^ melanoma cell lines. We evaluated the effect of GRPR activation on anoikis after seeding cells on poly-HEMA-coated plates with a low serum concentration and harvesting them 48 h later. Resistance to anoikis was assessed by the number of apoptotic cells (annexin-V positive) found under the various conditions. GRP induction promoted resistance to anoikis in mouse and human GRPR^pos^ cells (Figure 3J, 3K, 3N, and 3O) but not GRPR^neg^ melanoma cell lines (Figure 3L, 3M). Upon inhibition of GRPR with RC, the cells lost their resistance to anoikis (Figure 3J, 3K, 3N, and 3O). In addition, GRPR expression by itself was sufficient to significantly induce resistance to anoikis (Figure 3L-3O).

We also assessed the invasive capacity of melanoma cells on Matrigel^®^. GRP induction promoted GRPR^pos^ 1057 and MDA-MB-435S melanoma cell invasion, which was inhibited in the presence of RC (Figure 3P and 3Q). There was no effect on 1181 or 501-Mel GRPR^neg^ melanoma cells unless they were ectopically transfected with *Grpr* (Figure 3R-3U). These results were all confirmed using other GRPR^pos^ (1064 and 1014-Grpr) and GRPR^neg^ (1014) melanoma cell lines (Figure S3M-S3O). Note, that, in our conditions in the absence of GRP, the basal Grpr level was sufficient to significantly (p < 10^-4^) increase cell invasiveness (166 cells for [1181] *vs*. 367 cells for [1181-Grpr] (Figure 3R and 3T), 117 [501-Mel-Ctrl] *vs*. 360 [501-Mel-GRPR] (Figure 3S and 3U), 28 [1014-Ctrl] *vs*. 410 [1014-Grpr] (Figure S3N and S3O).

### GRPR signals through Gαq to activate YAP1 signaling

G protein-coupled receptors (GPCRs) signal through various Gα and downstream signaling pathways. We carried out a kinase assay using PamGene™ technology to identify the major pathway activated by GRP/GRPR in melanoma cells. GRPR activation was associated with PKC activation and, to a lesser extent, PKAα and PRKX activation (Figure 4A). PKCs are commonly activated by IP3/DAG signaling, secondary messengers of Gαq/11 signaling, whereas PKA is activated by the cAMP pathway downstream of Gαs. The activation of Gαq and PKC was supported by RNA-seq data after GRPR activation (Figure 4B). We evaluated activation of the receptor and its coupling with Gαq after measuring the production of IP3-IP1 after GRP induction. GRPR activation by GRP induced IP1 production in 1057 cells (GRPR^pos^), showing the activation of Gαq, and was inhibited in the presence of RC (Figure S4A). We observed no IP1 production by 1181-Ctrl cells (GRPR^neg^), but it was induced in 1181-Grpr cells (Figure S4A). Gαq has been shown to be associated with YAP1 activation in non-cutaneous melanoma models ^21, 22^. Moreover, all the cellular processes activated by GRPR can be explained by YAP1 activation ^3, 23, 24^. Indeed, the transcriptome of the mouse primary tumors showed that the YAP1 signature was induced in ΔEcad female tumors but not ΔEcad male nor Ecad melanomas (Figure S4B). Analysis of the TCGA transcriptome for human cutaneous melanoma showed a correlation between the level of GRPR and the YAP1 activation score (Figure 4C, r = 0.35). YAP1 was also found in the nucleus after Grp induction in 1057 cells (GRPR^pos^), but not in the presence of RC (Figure 4D). Transcriptomic analysis of 1057, 1181-Grpr, and 1014-Grpr cells (GRPR^pos^) showed the YAP1 activation signature to be significantly increased upon GRPR activation, but not in 1181-Ctrl and 1014-Ctrl (GRPR^neg^) cells (Figure 4E and S4C). In addition, the expression of YAP1 targets (CYR61, TEAD4, and LATS2) was induced after GRP stimulation, but repressed in the presence of GRP and RC, confirming the transcriptomic analyses (Figure 4F, S4E, and S4F). In conclusion, GRPR activation leads to Gαq activation which, in turn, stimulates the YAP1-regulated metastasis gene expression program.

**Figure 4.**
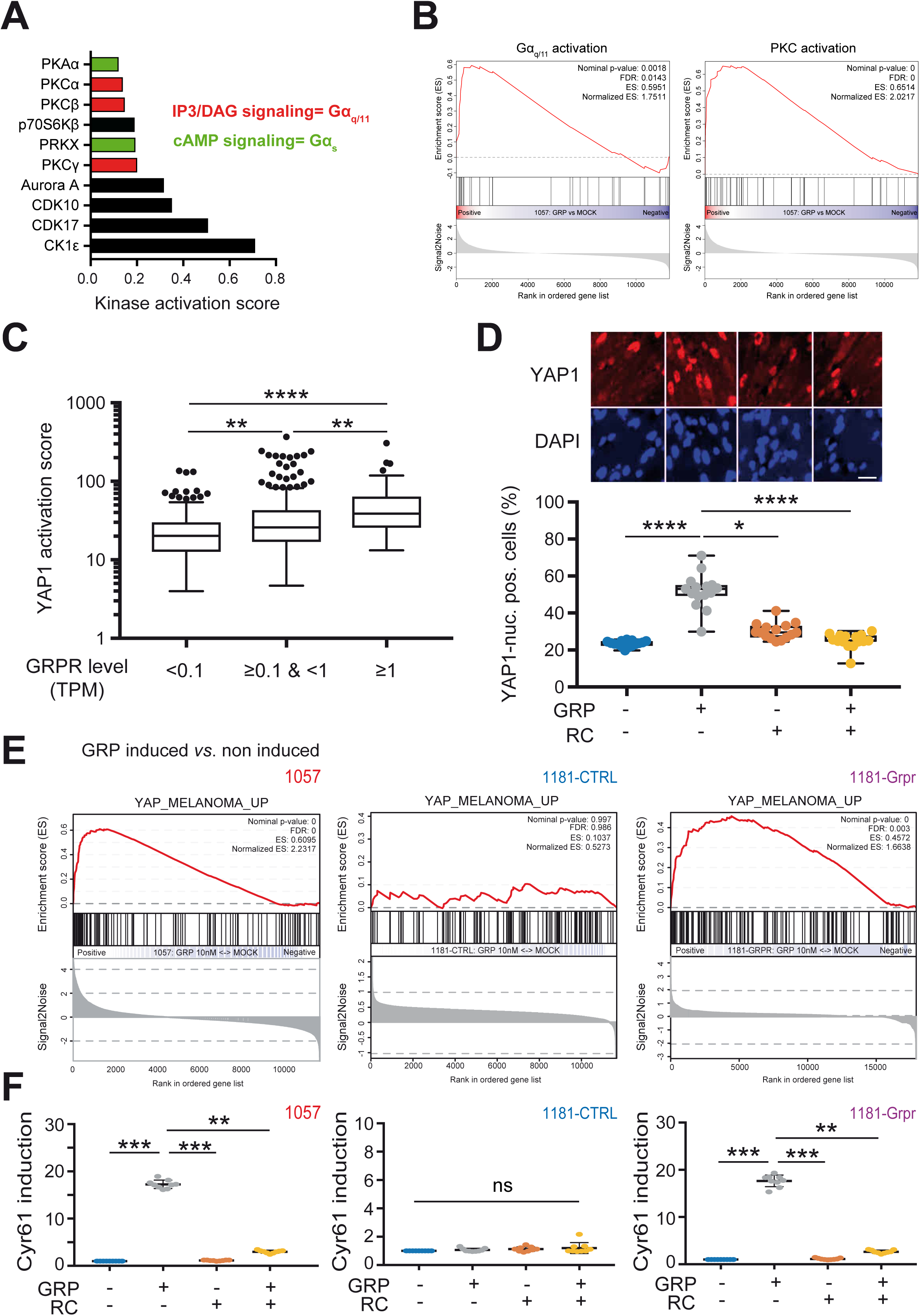
GRPR activation by GRP promotes the YAP1-transcriptional program. (**A**) Kinase activated by GRPR activation 15 min after induction by 10 nM GRP of the GRPR^pos^ melanoma cell line 1057. The kinase assay was conducted using the Pamgene® Ser/Thr kinase PamChip (STK). (**B**) Gene set enrichment analysis (GSEA) of the signatures of Gαq (left) and PKC (right) activation 4 h after the stimulation of 1057 cells (GRPR^pos^) with 10 nM GRP. Gene expression was measured by RNA-seq and read counts were normalized using DEseq2 prior to analysis. (**C**) YAP1 activation score according to the level of GRPR expression in melanomas from the TCGA-SKCM study. TPM: transcripts per million. (**D**) Representative pictures and percentage of cells with nuclear localization of YAP1 in the GRPR^pos^ mouse melanoma cell line (1057) after stimulation for 1 h with vehicle (lane 1), 10 nM GRP (lane 2), 1 µM RC (lane 3), or both (lane 4). Scale bar= 10 µm (**E**) GSEA of the YAP1 activation signature in GRPR^pos^ 1057 (left) and 1181-Grpr (right) and the GRPR^neg^ melanoma cell line 1181-Ctrl (middle). Gene expression was assessed by RNA-seq 4 h after stimulation with10 nM GRP and normalized using DEseq2 prior to conducting the GSEA. (**F**) Induction of the YAP1 target Cyr61 in the 1057 (left), 1181-Ctrl (middle), and 1181-Grpr (right) mouse melanoma cell lines. ns = not significant, *p < 0.05, **p < 0.01, ***p < 0.001, and ****p < 0.0001.

### GRPR activation promotes lung melanoma metastasis and can be targeted *in vivo*

The induction of proliferation, invasion, and resistance to anoikis by GRP/GRPR are key cellular processes for the colonization of distant organs by tumor cells. We assessed the capacity of GRPR to promote metastasis by injecting mouse 1181-parental, 1181-Ctrl (= Mock), and 1181-Grpr melanoma cells into the tail veins of immunocompetent C57BL/6J mice. Only GRPR-expressing cells (1181-Grpr) were able to colonize the lungs (Figure 5A and 5B). All mice injected with the 1181-Grpr cell line showed melanoma metastases in the lungs, with an average of 50 visible foci (Figure 5B and 5C). Similarly, human 501mel-parental, 501mel-Ctrl, and 501mel-GRPR cells were injected into the tail vein of immunodeficient (NSG) mice; 6 of 6 mice injected with 501mel-GRPR and 1 of 6 mice injected with 501mel-Ctrl showed lung metastasis (Figure 5D). Moreover, only one metastatic spot was found in the lung of this mouse injected with 501mel-Ctrl versus a median of 21 metastases in the 501mel-GRPR-injected mice (Figure 5E).

**Figure 5.**
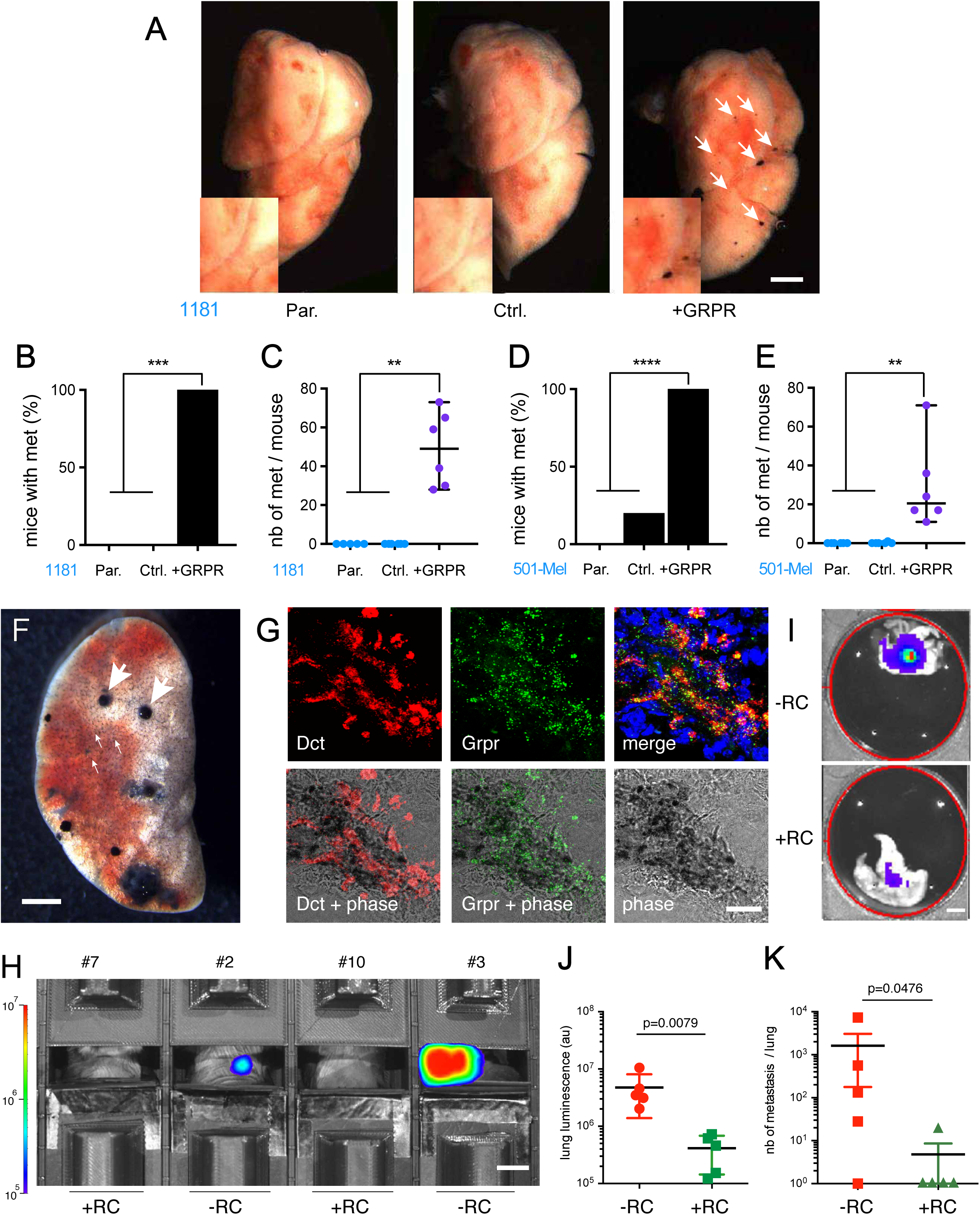
GRPR expression and activation induce lung melanoma metastasis. (**A**) Representative images of mouse lungs 30 days after the injection of 5.10^5^ 1181 melanoma cells not expressing GRPR (Par. and Ctrl.) or expressing GRPR exogenously (+GRPR) into the tail veins of C57BL/6J mice. Scale bar= 2 mm. (**B, C**) 1181 melanoma cells. **(D, E)** 5.10^5^ 501-Mel melanoma cells expressing GRPR (+GRPR) or not (Par. and Ctrl.) were injected into the tail veins of C57BL/6J NSG mice respectively. (**B, D**) Frequency of mice bearing lung melanoma metastases and (**C, E**) the number of lung metastases. Note, all 501mel-injected mice (parental, Ctrl, and GRPR) showed liver metastases but only the GRPR-expressing cells were able to efficiently colonize the lung. (**F-K**) Lung metastases after tail vein injection in C57BL/6J of 5 10^5^ 1057-Luc GRPR^pos^ cell RC-treated (mouse #6-10) or not (mouse #1-5). (**F**) Melanoma metastases present in the lung on day 28 after intravenous injection of 5.10^5^ 1057-Luc GRPR^pos^ melanoma cells. Note the entire colonization of the lung by the melanoma cells (small arrows) and proliferative foci (large arrows). Scale bar= 2 mm. (**G**) RNAscope of the lung of a 1057-Luc-injected mouse. Grpr (green) and Dct (red) mRNA colocalized in lung melanoma metastases. Scale bar= 50 µM. (**H**) Luminescence emerging from the thorax of mice after treatment or not with RC. Luminescence was acquired by IVIS. Scale bar= 1cm. (**I, J**) Evaluation of the luminescence of *ex vivo* lungs by IVIS after 28 days of treatment with RC or vehicle. Scale bar= 4 mm (**K**) Estimation of the metastases from five isolated independent lungs from mice treated or not with RC. Note that four lungs and one lung did not have any superficial metastases when they were treated or not with RC, respectively. Frequencies were compared using the Chi-square test and metastasis compared by one-way ANOVA or a two-tailed Mann-Whitney test when only two conditions were compared. Correction for multiple testing was performed using Tukey’s post-test. Data are presented as the mean values ± SD (**J**) or SEM (**L**). ns = not significant, *p < 0.05, **p < 0.01, ***p < 0.001, and ****p < 0.0001. Par. = parental.

We next performed a series of experiments to evaluate the potential of different drugs to inhibit the growth of GRPR-producing cells in the lungs in a longitudinal *in vivo* experiment. To visualize lung colonization by melanoma cells, we infected 1057 cells with a reporter pPGK-Luc2 construct to produce the 1057Luc melanoma cell line prior to tail-vein injection of female C57BL/6J mice. We observed numerous macro- and micro-lung metastases after the injection of 1057Luc cells. These results show the strong invasive potential and ability of these cells to proliferate in the lungs and colonize them (Figure 5F). Lung melanoma foci were pigmented and expressed both Dct and Grpr, as revealed by RNAscope experiments (Figure 5G). We tested the two antagonists of GRPR, RC-3095 (RC) and PD-176252 (PD) that had similar effects on 1057 cells (GRPR^pos^) growth *in vitro* (Figure S5A and S5B), for plasma and metabolic stability (Figure S5C and S5D). Despite the similar stability of the two drugs in plasma, PD was particularly sensitive to hepatic metabolic enzymes, with a half-life of 1.5 min versus 75 min for RC. We therefore used RC to evaluate *in vivo* lung colonization by 1057 cells (GRPR^pos^) by bioluminescent imaging (IVIS). Mice were injected in the tail vein and the success of injection evaluated by IVIS (Figure S5G). One day after tail-vein injection, mice were randomized into two groups (Figure S5E and S5F) and treated with either vehicle or RC. Luminescence in the thorax was first detected around day 24 and the signal increased exponentially (Figure S5G). We focused on signals originating from the lungs only by placing the mice in a custom box with shutters blocking out the signals of the thorax. Targeting GRPR with RC resulted in strong inhibition of lung colonization (Figure 5H-5K). On day 24, luminescence in the thorax was only detected in mice treated with vehicle but not those treated with RC (Figure 5H). The low level of lung colonization by 1057Luc cells when treated with RC was confirmed by *ex vivo* luminescence of the lungs (Figure 5I, 5J, and S5H). Luminescence values were consistent with the number of visible lung metastasis (Figure 5J and 5K). In conclusion, treatment of the mice with RC strongly impaired colonization of the lungs by GRPR-expressing melanoma cells.

### GRPR is expressed in females through a CDH1-ESR1-GRPR axis

The female-specific activation of Grpr expression led us to analyze the pathways regulated by sex hormones. Comparison of the transcriptome of the Ecad and ΔEcad female tumors by GSEA analysis showed the ΔEcad tumors to overexpress genes associated with activation of the estrogen receptor alpha (ERα) (Figure 6A). This signature was associated with the expression of ERα, encoded by the ESR1 gene, in the ΔEcad female melanoma cell lines, suggesting that the regulation of Grpr by estrogen is cell autonomous (Figure 6B). None of the other sexual hormone receptors (*Esr2*, *Gper1*, *Ar*, and *Pgr*) were expressed in the murine melanoma cell lines. Silencing the expression of Ecad in murine and human melanoma cells led to an increase in Grpr mRNA expression, supporting a cell-autonomous mechanism (Figure 6C). This supports the anti-correlation between CDH1 and GRPR expression observed in the TCGA data (Figure 2N). Silencing of ESR1 in murine and human ESR1^pos^/GRPR^pos^ melanoma cells induced a significant decrease in GRPR mRNA levels (Figure 6D), whereas ectopic ESR1 expression increased GRPR mRNA levels (Figure 6E). Ecad silencing, in addition to increasing GRPR expression, induced a significant increase in ESR1 mRNA expression, confirming the inhibition induced by Ecad on Esr1 mRNA expression (Figure 6F). Silencing of both CDH1 and ESR1 in murine and human melanoma cell lines blocked the upregulation of GRPR expression, showing that its following Ecad loss is due to the induction of ESR1 expression (Figure 6F). The CDH1/ESR1/GRPR axis was regulated by a positive feedback loop, as GRPR activation repressed the expression of CDH1 (Figure 6G). Estrogens fuel this loop, as β-estradiol, an ERα agonist, significantly increased GRPR expression and treatment with ICI 182,780, an ERα antagonist, repressed GRPR expression (Figure 6H). Thus, following the loss of E-cadherin expression, a transcriptional regulatory program is initiated and amplified by a positive regulatory loop, explaining the high expression of *GRPR* (Figure 6I).

**Figure 6.**
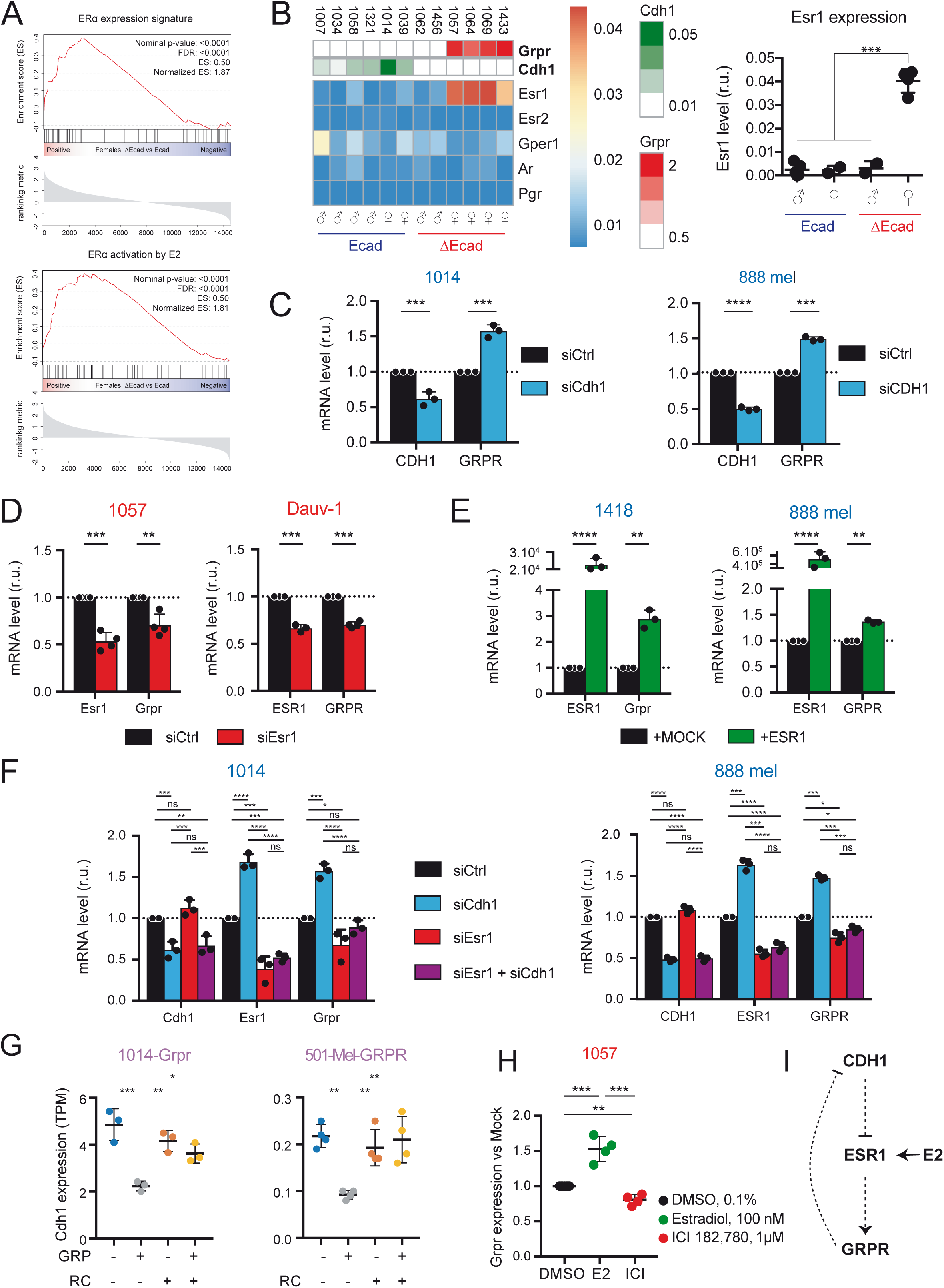
GRPR is expressed in females through a CDH1/ESR1/GRPR amplification loop. (**A**) GSEA of the ERα expression signature (top) and ERα activation by E2 estrogen (bottom) comparing murine ΔEcad female primary melanoma to Ecad female primary melanoma. (**B**) Heatmap depicting the relative expression of sexual hormone receptors in Ecad and ΔEcad murine melanoma cell lines of both sexes annotated with the corresponding Cdh1 and Grpr expression (at the top). All expression was evaluated by qRT-PCR and normalized to Hprt expression. The right panel shows Esr1 expression in various cell lines. (**C**) Effect of the knock-down of *CDH1* expression on *GRPR* levels in mouse (1014, left panel) and human (888-Mel, right panel) CDH1^pos^ melanoma cell lines. (**D**) Effect of the knock-down of *ESR1* expression on *GRPR* expression in murine (1057, left) and human (Dauv-1, right) ESR1^pos^ melanoma cell lines. Expression was assessed 48 h after cell transfection. (**E**) Effect of the overexpression of *ESR1* on *GRPR* expression in murine 1418 (left) and human 888 Mel (right) ESR1^neg^/GRPR^neg^ melanoma cell lines. Expression was assessed 48 h after cell transfection. (**F**) Consequences of the silencing of *CDH1* and/or *ESR1* expression on GRPR expression in CDH1^pos^ mouse melanoma cells (1014 – left panel) and human melanoma cells (888 Mel – right panel). Expression was assessed 48 h after cell transfection. (**G**) Impact of GRPR activation on CDH1 expression in murine 1014 (left) and human 510-Mel CDH1^pos^ melanoma cell lines with exogenous expression of GRPR. Cells were stimulated with 10 nM GRP and/or treated with 1 µM RC for 4 h. Expression was assessed by RNA-seq followed by normalization to the TPM level. (**H**) Consequence of the activation of ERα by 100 nM β-estradiol (E2) or the degradation of ERα by 1 µM ICI 182,780 on *GRPR* expression 48 h after treatment in ESR1^pos^ 1057 melanoma cells. Cells were starved of estrogen and phenol red for four days prior to treatment. (**I**) Scheme summarizing the results. CDH1 represses *ESR1* expression. The loss of CDH1 in melanomas induces the expression of *ESR1*. ERα, encoded by *ESR1*, promotes *GRPR* expression. Finally, GRPR activation decreases *CDH1* expression, resulting in an amplification loop. All comparisons were performed by ANOVA. Adjustments for multiple comparisons were performed using Tukey’s post-test. ns = not significant, *p < 0.05, **p < 0.01, ***p < 0.001, and ****p < 0.0001.

### The CDH1-ESR1-GRPR axis is active in breast carcinomas

As GRPR and CDH1 expression are anticorrelated in breast tumors (Figure 2O) and the growth of many breast-tumor cells is estrogen dependent ^25^, we hypothesized that the repression of the ESR1/GRPR axis by E-cadherin is fundamental and occurs in other tumors, such as breast carcinomas. The anticorrelation was supported by significantly higher GRPR mRNA levels in tumors expressing a mutated form of non-functional Ecad (Figure 7A). As in melanoma, the silencing of CDH1 increased GRPR expression in MCF7 breast cancer cells (Figure 7B). The dependence of GRPR expression on ERα levels was supported by the higher levels of GRPR in ERα-positive breast tumors than in ERα-negative breast tumors (Figure 7C) and by the strong correlation (r = 0.52, with a highly significant p-value) of GRPR expression with the ERα activation score (Figure 7D). Silencing of ESR1 expression in MCF7 cells resulted in a significant decrease in the expression of GRPR, confirming the regulation of GRPR by ERα (Figure 7E). The activation of ERα by estradiol resulted in an increase in GRPR expression, whereas the drug ICI 182,780 substantially reduced GRPR levels (Figure 7F). BRCA-tumors mutated for *CDH1* expressed higher levels of ESR1 and had a higher ERα activation score and higher positivity of ERα expression, as assessed by IHC (61.2% in *CDH1* WT tumors vs 94.5% in *CDH1* mutated tumors) (Figure 7G and 7H). Overall, these data confirm that the silencing of CDH1 increases both ESR1 and GRPR expression, whereas the concomitant silencing of CDH1 and ESR1 does not induce GRPR expression in MCF7 cells (Figure 7I).

**Figure 7.**
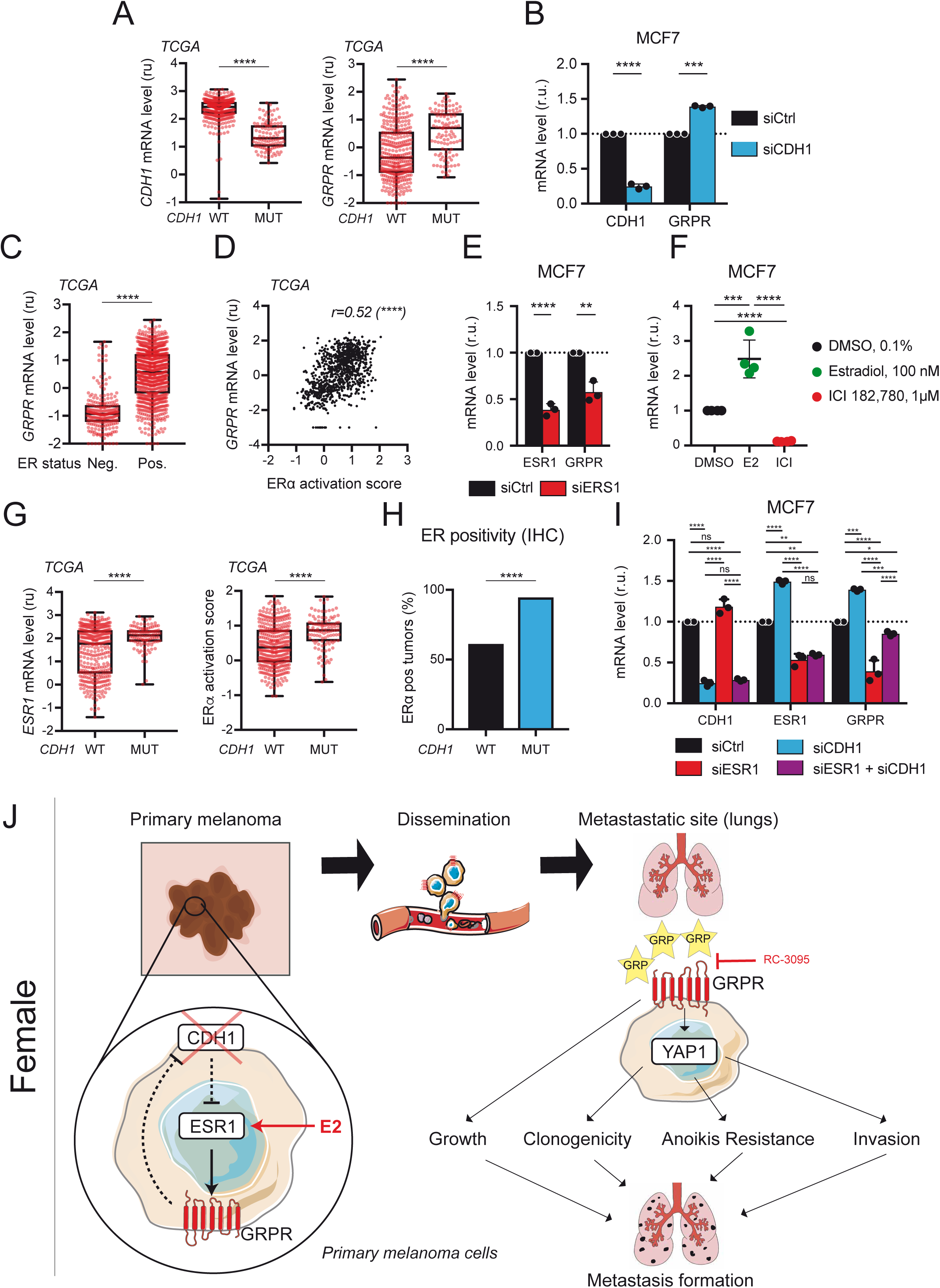
The CDH1-ESR1-GRPR axis is also active in breast carcinomas. (**A**) Expression of CDH1 (left) and GRPR (right) mRNA in breast carcinoma from TCGA-BRCA data according to the *CDH1* genetic status. (**B**) Consequence of CDH1 knock-down on CDH1 and GRPR expression in the MCF7 breast cancer cell line. (**C**) Expression of GRPR according to the ERα status defined by the pathologist by immunohistochemistry (IHC) in human breast carcinomas from the TCGA-BRCA studies. (**D**) Correlation between GRPR expression and the ER □ activity score calculated from the same breast tumor originating from the TCGA-BRCA studies. The significance of the correlation was assessed using the Pearson test. (**E**) Effect of ESR1 knockdown on ESR1 and GRPR expression in the MCF7 breast cancer cell line. (**F**) Effect of ESR1 stimulation or inhibition on GRPR expression. MCF7 cells were starved of phenol-red and estrogen for four days and then stimulated for 48 h in phenol-red free serum-stripped media containing DMSO (0.1%, left lane), the ERα agonist 17β-estradiol (100 nM, center lane), or the ERα antagonist ICI 182,780 (1 µM, right lane). (**G**) Expression of ESR1 mRNA in breast cancer (left) and the ER activation score (right) according to the genetic status of *CDH1* in breast cancer tumors from the TCGA data. (**H**) Quantification of ERα positivity of breast carcinomas from the TCGA-BRCA cohort by IHC according to *CDH1* mutation. (**I**). Consequence of the silencing of *CDH1* and/or *ESR1* expression on GRPR expression in the CDH1^pos^ MCF7 human breast cancer cell line. Expression was assessed 48 h after cell transfection. Comparison of the siRNA effects between two groups was performed using a multiple t-test corrected by the FDR method. Comparison of the siRNA effects between more than two groups was performed by ANOVA and adjusted for multiple comparisons using Tukey’s post-test. Other comparisons were performed using a two-way t-test, except for the comparison of proportions, which was performed using a two-way Chi-square test. ns = not significant, *p < 0.05, **p < 0.01, ***p < 0.001, and ****p < 0.0001. The relative mRNA levels in **A, C, D,** and **G** correspond to the Log10 (TPM + 0.01). (**J**) In the primary melanoma, the loss of CDH1 induces the expression of *ESR1*. In turn ERα activates the expression of *GRPR,* which is stronger in the presence of estrogen (E2) especially between puberty and menopause. GRPR activation by GRP decreases *CDH1* expression, amplifying this loop and leading to high levels of GRPR expression (left part). The Ecad^neg^/GRPR^pos^ melanoma cells are able to disseminate through the vasculature to distant organs, especially the lungs, where GRP is abundantly produced. In the lungs, the interaction between GRP/GRPR induces pro-metastatic signaling in the melanoma cells through the activation of YAP1. The activation of key cellular mechanisms involved in melanoma progression, such as cell growth, cell survival, cell clonogenicity, resistance to anoikis, and cell invasion, ultimately lead to the formation of metastases in the lungs. This mechanism can be impaired *in vivo* by targeting GRPR with an antagonist such as RC (right part).

## Discussion

In the present study, we describe a metastatic pathway induced by the loss of Ecad that is specifically activated in women. Reduction in the level of ECAD induces an increase in the level of the estrogen receptor ESR1, which in turn induces the transcription of a G-protein coupled receptor, GRPR. Activation of GRPR induces activation of YAP1, which leads to metastasis through the induction of key cellular processes in cancer progression, including proliferation, migration/invasion, and resistance to anoikis. This CDH1-ESR1-GRPR-YAP1 axis defines a sex-specific metastatic pathway in tumors of females that opens new therapeutic opportunities (Figure 7J).

Our study shows that the level of GRPR markedly increases after the loss of ECAD in women. This result shows that ECAD acts as a tumor suppressor by repressing GRPR in melanoma. In breast cancer cells, repression of GRPR by ECAD is also observed. An anti-correlation between CDH1 and GRPR has been observed in many carcinomas, indicating that this is a general regulatory mechanism. Interestingly, the repression of GRPR by ECAD has been notably observed in two carcinomas, those of the breast and stomach, in which ECAD has been previously shown to be a tumor suppressor ^14, 26^. Furthermore, GRP, the natural agonist of GRPR, is abundantly produced in the breast and stomach ^27^. GRPR is likely activated in the primary tumor, whereas in other carcinomas or in melanoma, GRPR activation likely only occurs at the metastatic site, where GRP is produced. We propose that ECAD acts as a suppressor of tumor initiation when the primary tissue produces GRP and acts as a suppressor of tumor metastasis when GRP is not produced in primary tissue but at metastatic sites, such as the lung for cutaneous melanoma.

Recently, there has been a growing awareness of the need to better account for parameters outside of the disease, such as ethnicity, age, sex, or gender in the management of cancer patients, as large disparities in the incidence and response to therapy have been clearly identified ^28–32^. For example, the incidence of melanoma is higher in women between puberty and menopause ^33, 34^ and men respond better to immunotherapies, whereas women respond better to MAPKi-targeted therapies ^35, 36^. These data support the hypothesis that hormones are important in the development and treatment of cancer. By placing ERα as a major player in the progression of melanoma, as well as that of carcinomas, our study provides clear and complementary evidence that estrogens are key micro-environmental factors in carcinogenesis. ERα can be activated by mutation in 1.4% of melanoma cases, according to the TCGA database, but more classically by the binding of estrogens produced in abundance by women between puberty and menopause. ERα can also be activated in men and women by xenobiotics, including estrogenic contraceptives, pesticides, and plastic derivatives. The CDH1-ESR1-GRPR activation loop that we present can be initiated at different levels: (i) alterations of Ecad by mutations, methylation, and/or EMT ^16, 26, 37^, (ii) activation of ER by the menstrual cycle, pregnancy, and/or xenobiotic, mutations ^38, 39^, and (iii) activation of GRPR by GRP, other endogenous low-affinity ligands, and/or transactivation by interaction with other receptors ^40^.

We show that Ecad represses ESR1 expression in melanoma and breast cancer cells in a sex-dependent manner in both humans and mice. The exact mechanisms that link regulation of the ESR1 gene by ECAD are still unknown, although we previously reported that ECAD regulates gene expression ^10^. Cadherins can regulate gene expression after interacting with various partners, including receptor tyrosine kinases, such as EGFR and IGFR for E-cadherin ^41, 42^, FGFR for N-cadherin ^43^, and VEGFR for VE-cadherin ^44^. These interactions inhibit the activity of the receptor tyrosine kinases with which they interact and which are known to regulate specific signaling pathways ^45, 46^. Cadherins can also regulate gene expression after interacting with catenins, such as β-catenin, plakoglobin (= catenin), p120-catenin, and/or α-catenin. The lack of ECAD allows the release of these proteins from the membrane and then may induce their signaling. However, in our conditions, activation mediated by catenins was not observed. Similar results were previously shown in ES cells ^10^. In addition, Ecad can modulate the chromatin landscape, as shown in an EMT model in which low Ecad states modulate CTCF expression and consequently chromatin conformation ^47^. A comprehensive and thorough study of chromatin conformation is needed to better understand how a cell-cell adhesion molecule can shape chromatin architecture.

The identification of the CDH1-ESR1-GRPR-YAP1 axis has led to the recognition of three potential therapeutic targets (ERα, GRPR, and YAP1) to treat cancer directly or in combination with current therapies. Modifying ERα activity in breast cancer by targeting the receptor directly with antagonists or by targeting estrogen synthesis decreases the growth and/or progression of this cancer ^48–50^. Because ERα activity correlates with GRPR expression, we hypothesize that down-regulation of GRPR could explain this therapeutic action. This strategy could thus be applied to other cancers, notably to hinder the formation of lung melanoma metastases in women. GPCRs are widely targeted, but their targeting is still underutilized in cancer ^51, 52^. Here, we identified GRPR as a major player in cancer progression. We have demonstrated that targeting GRPR with RC-3095 in melanoma decreases the growth of lung metastases. Moreover, it has been suggested that this antagonist reduces the growth of breast, lung, prostate, and pancreas tumor cell lines *in vitro* and *in vivo* ^53–56^. However, drugs that target GRPR are in great need of improvement, either because their pharmacokinetic properties are not sufficient for effective use *in vivo* or because of excessive toxicity ^57^. The generation of new selective GRPR antagonists is therefore critical. Moreover, antagonists that are specific should result in limited side effects, Grpr is weakly expressed in adults and *Grpr* knockout mice do not show a severe phenotype, apart from satiety and pruritus. ^58, 59^. Finally, YAP1 has been shown to be associated with cancer progression and our study confirms that targeting YAP1 in this context could indeed be effective ^60, 61^. Although targeting YAP1 may be effective for cancers that are not GRPR positive, its safety profile is likely to be lower than that of GRPR antagonists.

In conclusion, this study highlights the importance of considering gender diversity in human pathologies as a crucial factor in the therapeutic management of patients. Our results provide a novel gender-specific therapeutic lead to improve cancer treatment in women, in whom the loss of E-cadherin results in ERα and GRPR expression.

## Acknowledgments

We are grateful to Dr. Daniel Metzger for providing ESR1 expression vector and for helpful comments. We thank Drs. Martin Dutertre and Delphine Duteil for insightful conversations. We thank Drs. Beermann, Kemler and Seano for providing Tg(Tyr-NRAS*Q61K)1Bee, B6.129-Cdh1tm2Kem/J and Cdkn2a^tm1Rdp^ mouse lines respectively. We thank John Groten, Mayke Bel, Eddy Van Breemen, Savithri Rangarajan, and Justin lebens from Pamgene®. We thank the Institut Curie staff responsible for the animal colony (especially P. Dubreuil and C. Lantoine), the histology (S. Leboucher), FACS (C. Lasgi), and PICT-IBiSA imaging (C. Lovo and L. Besse) facilities. This work was supported by FRM EQU202103012599, Institut Curie, INSERM, and CNRS. J.R. had a fellowship from PSL and LNCC. M.W. had a fellowship from DIM SEnT. F.L. had a fellowship from MENRT, SFD and LNCC.

## Author contribution

L.L. and V.D. conceived and supervised the study. J.H.R, M.S., V.P., Z.A., F.L., M.W., P.G., C.B., B.V. and V.D. performed experiments and J.H.R, M.S., V.P., Z.A., F.L., M.W., B.V. L.L. and V.D. analysed data. C.L.T. and F.M.B. provide critical resources. IM provides expertise. J.H.R., L.L. and V.D. produced the graphical representation of the data. L.L. and V.D. acquired funding to support this project. The manuscript was written by J.H.R., L.L. and V.D. and reviewed by V.P., Z.A., I.D. and F.M.B. All authors approved of and contributed to the final manuscript.

## Inclusion and diversity

We support inclusive, diverse, and equitable conduct of research.

## STAR Method

### KEY RESOURCES TABLE

See attached word file

### RESSOURCE AVAILABILITY

#### Lead contact

Further information and requests for resources and reagents should be directed to and will be fulfilled by the lead contact, Lionel LARUE.

#### Materials availability

All newly generated materials or mouse lines used in this study are available upon request to the lead contact with a completed Materials Transfer Agreement.

#### Data code and availability

RNA-seq data have been deposited at GEO and are publicly available as of the date of publication. Accession numbers are listed in the key resources table. Microscopy data reported in this paper will be shared by the lead contact upon request.

This paper does not report original code.

Any additional information required to reanalyze the data reported in this paper is available from the lead contact upon request.

## EXPERIMENTAL MODEL AND SUBJECT DETAILS

### Mice

Animal care, use, and all experimental procedures were conducted in accordance with recommendations of the European Community (86/609/EEC) and Union (2010/63/UE) and the French National Committee (87/848). Animal care and use were approved by the ethics committee of the Curie Institute in compliance with institutional guidelines. Experimental procedures were carried out under the approval of the ethics committee of the Institut Curie CEEA-IC #118 (CEEA-IC 2016-001) in compliance with international guidelines. The transgenic *Tyr::CreA* (B6.Cg-Tg(Tyr-cre)1Lru/J), named Tyr::Cre, Tg(Tyr-NRAS*Q61K)1Bee, Cdkn2a^tm1Rdp^, and B6.129-Cdh1tm2Kem/J, named Cdh1F/F, mice have been described and characterized previously in the Larue laboratory and elsewhere ^18, 19, 62^. The mouse lines were backcrossed onto a C57BL/6J background for more than 10 generations. Genotyping was performed according to ^63^ using specific primers and conditions (Table S6-S7). Mice were crossed to obtain the desired genotypes. Mice were born with the expected ratio of Mendelian inheritance and no changes in gender ratios were observed. Mice were checked weekly for the appearance of new tumors. Tumors were allowed to grow until reaching a volume of 1 cm^3^ and then autopsied to analyze the presence of metastases in distant organs.

### Tail-vein injection and IVIS follow-up

Cells (5 10^5^) of mouse and human melanoma cell lines were injected in 200 µL PBS into the tail veins of eight-week-old C57BL/6J (RRID:IMSR_JAX:000664) and NSG (RRID:IMSR_JAX:005557) mice respectively. For each experiment, six mice per group were injected. Mice were followed daily and weighed twice a week. Mice were euthanized and autopsied when they had lost 20% of their maximal weight. For IVIS follow up, C57BL/6J mice were shaved three days before cell injection and then every other week. Five mice per group were injected with cells of the 1057 melanoma cell lines and then treated. Briefly, mice were injected with 300 µg Xenolight D-luciferin and immediately anesthetized with isoflurane. Ten minutes later, they were placed in an opaque box with adaptable shutters to image only the thorax in the IVIS apparatus (IVIS spectrum, #124262, Perkin-Elmer). Luminescence from the whole mouse and the thorax alone were acquired successively for 2 min each. Luminescence was acquired on day 0, day 1, and then twice a week. The day after cell injection, mice were randomized into two groups according to their weight and luminescence in the lung at day 0. Each group of mice was treated twice a day with either 10 µg RC/DMSO or vehicle PBS/DMSO for one month and followed by IVIS. After euthanasia, the lungs were carefully removed from the thorax and quickly rinsed in PBS. The right lungs were incubated in a black opaque 12 well plate with 300µg/mL Xenolight D-luciferin for 2 min and luminescence acquired for 1 min. The left lungs were fixed for 24 h in 4% PFA at 4°C and incubated in 30% sucrose and a 30% sucrose/50% OCT solution for 48 h each and then embedded in OCT. OCT-embedded mouse lungs were sectioned into 7-μm-thick coronal sections. Sections were stained for Dct and Grpr mRNA and counterstained using the manufacturer’s protocol (RNAscope). RNAscope Mm-Dct-C1 (#460461, ACD) and Mm-Grpr-C3 (#317871-C3, ACD) probes were purchased from BioTechne. Images were acquired using an inverted SP8 Leica confocal microscope (Leica Microsystem). Right lungs were imaged using a Leica MZFLIII binocular microscope equipped with a Scion camera. Metastases were enumerated using ImageJ. Macro-metastases were defined by a diameter > 0.1 mm and micro-metastasis < 0.1 mm.

### Cell lines

Mouse melanoma cell lines were established from melanomas arising in transgenic mice in the laboratory as previously described ^64^. MDA-MB-435S, 501-Mel, 888 Mel, and Dauv-1 human melanoma cell lines were previously established in other laboratories ^65–67^. The human breast cancer cell line MCF-7 was previously established ^68^. The pGK-Luc2 vector was a gift from Catherine Tomasetto (IGBMC, Strasbourg, France). Briefly, the coding sequence of the luciferase reporter gene luc2 (Photinus pyralis) was amplified by PCR from the pGL4.50[luc2/CMV/Hygro] vector (Promega E1310) and flanking XhoI restriction sites were added. The digested PCR fragment was subcloned into the SalI site of the pLENTI PGK Blast DEST vector (Plasmid #19065, Addgen). 1057-luciferase melanoma cell lines were generated by infecting parental 1057 cells with pGK-Luc2 (LL#1231). Cells were selected using 4.5 µg/mL Blasticidin for one week. Lines 1014 and 1181 Grpr and the corresponding controls were obtained after transfection of the murine Grpr/tGFP plasmid (#1045 MG224721, Origene) and tGFP plasmid (#1064 pCMV6-AC-GFP, Origene), respectively. Cells were transfected with 2 to 4 µg of plasmid or 100 pmol of siRNA and lipofectamine 2000 following the manufacturer’s protocol. Transfected 1014 and 1181 cells were selected using 25 and 150 µg/mL geneticin, respectively. 501-Mel-GRPR and 501-Mel-Ctrl were generated by infecting cells with the pLV-Hygro-CMV-GRPR-EGFP (#1VB191126-1286xxe, LL1276, Vector Builder) and pLV-Hygro-CMV-EGFP (#VB191126-1289cfv, LL1271b, Vector Builder) plasmids, respectively. All siRNA used are listed in Supplementary Table S8.

Murine and human melanoma cell lines were grown in Ham’s F12 medium and RPMI 1640, respectively, supplemented with 10% FCS and 1% PS. The breast cancer cell line MCF7 was grown in DMEM-F12 supplemented with 10% FCS and 1% PS. All cell lines were maintained at 37°C in a humidified atmosphere with 5% CO2. Cells cultured without phenol-red were supplemented with 2 nM glutamine. The genetic status and level of expression of key genes of these cell lines are presented in Table S5.

### Human samples

The retrospective study on lung human melanoma metastases was approved by the ethics committee. The non-opposition or consent (before or after 2004, respectively) of patients for the use of their biological material and data was obtained according to the bioethics law of 2004. Forty-three tissue samples of non-treated lung melanoma metastases (LMM) registered from 1999 to 2014 were retrieved from the pathology files of the Bordeaux and Rennes Hospitals. We selected all available FFPE surgical specimens of LMM for further immunostaining analysis.

## METHOD DETAILS

### TCGA data mining

The transcriptome, copy number alteration, mutations, and corresponding clinical data of the TCGA-SKCM (n = 473), TCGA-BRCA (n=1215), TCGA-STAD(n=450), TCGA-LUSC (n=553), and TCGA-KIRC (n=606) data sets were retrieved from the National Cancer Institute (NCI) Genomic Data Commons (GDC) repository using the TCGAbiolinks R package in August 2022 ^69^. mRNA levels were calculated from RNA sequencing read counts using RNA-Seq V2 RSEM and normalized to transcripts per million reads (TPM). Survival analyses were carried out by separating the cohort into two groups based on gene expression. The threshold was set to 1 TPM, commonly considered to be the limit for sufficient protein expression of the transcript. The negative group was set to an expression ≤ 0.1 TPM to have clear separation in terms of expression (factor 10) from the positive group. The YAP and melanoma phenotypic state scores were obtained by averaging the expression of detailed marker genes or for the mice, from their murine orthologs. The pigmentation state was defined by the expression of MITF, MLANA, TRPM1, DCT, and TYR; the SMC phenotype by the expression of CD36, DLX5, IP6K3, PAX3, and TRIM67; the invasive state by the expression of AXL, CYR61, TCF4, LOXL2, TNC, and WNT5A, and the NCSC-like phenotypic state by the expression of AQP1, GFRA2, L1XAM, NGFR, SLC22A17, and TMEM176B. The scoring of YAP1 activation was determined from the expression of CYR61, CTGF, TEAD4, LATS2, and CRIM1. Yap scoring was obtained by averaging the fold change of each Yap1 target. The anoikis-resistance score was calculated based on the expression of S100A7A, MTPN, ATP10B, S100A8, RSAD2, RENBP, CDHR1, and CD36. The ER-activation score was calculated based on the expression of GATA4, SDK2, EGR3, IL19, GSG1L, RSPO1, PGR, IL24, and PDZK1.

### Immunohistochemistry

Paraffin was melted at 56°C overnight. Deparaffinization, using a Bond Dewax Solution (Cat#AR9222, Leica), and rehydration were performed with a Leica BONDTM -MAX device. Heat-induced epitope antigen retrieval was performed at 100°C for 20 min in Bond Epitope Retrieval Solution 1 (Cat#AR9961, Leica) for GRPR or Bond Epitope Retrieval Solution 2 (1/100, Cat# AR9640, Leica) for Ecad. Slides were incubated in anti-GRPR antibody (SP4337P, Acris Antibodies) in Bond Primary Antibody Diluent (1/100, Cat# AR9352, Leica) overnight at 4°C and anti-Ecad (1/100, Cat#NCL-L-E-Cad, Novocastra) antibody in the same diluent for 30 min at room temperature (RT). Bond Polymer Refine Red Detection (Cat#DS9390, Leica) was used according to the manufacturer’s specifications. Slides were counterstained with hematoxylin and cover slipped. Images were acquired using an Axio Imager Z2 microscope. Each immunostaining was evaluated in a double-blind manner by two pathologists.

### Cell growth and clonogenic assay

Six-well tissue culture plates were seeded with 3.10^5^ melanoma cells in complete medium. Twenty-four hours later, the medium was replaced by low-serum medium (0.5% for murine melanoma cell lines and 1% for human melanoma cells) and the cells incubated a further 18h before stimulation with 10 nM GRP and/or 1 µM RC for 48 h. The plates were trypsinized just after stimulation or 48 h later and the cells were counted. Cell growth was assessed by MTT assay. Briefly, 10,000 cells were seeded per well in 96-well plates. Twenty-four hours later, cells were starved overnight and then treated with 10 nM GRP and/or 1 µM RC for 48 h in low-serum medium (0.5% FBS). Next, MTT was added to the wells to a final concentration of 0.5 mg/mL and the plates incubated for 3 h. The medium was removed and formazan crystals dissolved in 200 µL DMSO. Absorbance was read at 570 nM using a LUMIstar Omega luminometer (BMG Labtech). All growth experiments were conducted using three technical replicates and three biological replicates. For the clonogenic assay, six-well tissue culture plates were seeded with 500 cells in complete medium. After 10 days (20 days for 1181), colonies were fixed with 4% PFA for 15 min and stained with 10% crystal violet in ethanol for 20 min and counted in images using ImageJ software. Experiments were performed in triplicate.

### Matrigel invasion assay

Matrigel invasion assays were performed in transwell plates with 8.0 µm pores (#353097, Falcon) coated with 100 µL of 200µg/mL Matrigel. Cells were seeded in low serum medium (0.5% FCS for murine cells and 1% for human cell lines) containing 10 nM GRP and/or 1 µM RC as attractant or 10% FCS. Twenty-four hours after stimulation, inserts were washed with PBS and non-invading cells removed. Cells in the inserts were fixed in methanol at -20°C overnight. The inserts were rinsed and the membrane carefully removed using a sharp scalpel. The membrane was mounted in prolong glass DAPI (1.5 µg/mL). Assays were performed in triplicate and the entire filter was counted using Image J software.

### Anoikis assay

Six-well plates were coated with poly-HEMA to avoid cell attachment to the well surface. Cells were seeded in low-serum medium containing 10 nM GRP and/or 1 µM RC. Forty-eight hours after cell seeding, cells were harvested and washed with ice-cold PBS twice and resuspended in annexin V binding buffer (#556454 BS bioscience) and incubated at RT in the dark with 7-amino-actinomycin D (7-AAD, #559925, BD bioscience) and annexin-V for 15 min. Annexin V was coupled to FITC (#556420, BS bioscience) for the 1057 and MDA-MB-435S cell lines and to PE (#556421; BD bioscience) for the other cell lines. Cells were sorted using a FACS LSRFortessa (BD Bioscience) to determine the percentage of annexin V- and/or 7-AAD-positive cells using a 488 nm laser for annexin V-FITC and annexin V-PE and a 675 nm laser for7-AAD. All quantification was performed using Flow-Jo.

### Kinase assay

Serine/threonine and tyrosine kinase activity were determined using STK PamChips (#87102 PamGene International B.V.). All assays were performed according to the manufacturer’s protocol ^70^. Briefly, cells were seeded to 70% confluence and starved overnight the day after. Then, the medium was replaced by medium containing vehicle (0.1% DMSO) or 10 nM GRP and/or 1 µM RC and the cells incubated for 15 min. Cells were rinsed twice in PBS and then lysed in M-Per buffer (#78503 Thermo-Fisher Scientific) containing Halt Phosphatase Inhibitor (#78428 Thermo-Fisher Scientific) and Halt Protease Inhibitor (#78437 Thermo-Fisher Scientific), both diluted 1:100. The lysates were immediately snap-frozen. Kinase activity was determined using the Pamstation PS12 and 1 µg protein for the STK chips according to the manufacturer’s protocol. The data were analyzed using BioNavigator software (PamGene International B.V.), batch corrected using ComBat ^71^, and normalized using VSN ^72^. Kinase activity was assessed using the 2018 version of the UKA tool using basic parameters (Scan rank from 4 to 12, 500 permutations, 90% homology, equivalent weight for each database, minimal prediction score of 300). The UKA tool infers the active kinase from the differentially phosphorylated peptides using databases and predicted interactions (PhosphoNet database).

### RNA extraction and transcriptomic analysis

RNA was extracted from cells and mouse tumors using the miRNeasy kit (Qiagen, #217004) according to the manufacturer’s protocol. RNA integrity (RIN) was measured using an Agilent Bioanalyser 2100 (Agilent Technologies, Les Ulis, France). Only RNA with an RNA integrity number (RIN) > 7 was used for analysis. This threshold led to the sequencing of 72 mouse melanoma cell lines, 12 human melanoma cell lines, and 32 mouse tumors. RNA concentrations were measured using a NanoDrop (NanoDrop Technologies, Wilmington, DA, USA). RNA sequencing libraries were prepared from 1 μg total RNA using the Illumina TruSeq Stranded mRNA Library preparation kit, which allows strand-specific sequencing. PolyA selection using magnetic beads was performed to focus the sequencing on polyadenylated transcripts. After fragmentation, cDNA synthesis was performed and the resulting fragments used for dA-tailing, followed by ligation with TruSeq indexed adapters. The fragments were the amplified by PCR to generate the final barcoded cDNA libraries (12 amplification cycles). The 12 and 32 libraries were equimolarly pooled and subjected to qPCR quantification using the KAPA library quantification kit (Roche). Sequencing was carried out on a NovaSeq 6000 instrument (Illumina) based on a 2×100 cycle mode (paired-end reads, 100 bases) using an S1 flow cell to obtain approximately 35 million clusters (70 million raw paired-end reads) per sample. Reads were mapped to the mm10 mouse reference genome (gencode m13 version-GRCm38.p5) or hg38 human reference genome (gencode 42 version-GRCh38.p13) using STAR ^73^. STAR was also used to create the expression matrices. When applicable, expression was batch-corrected with Combat using the sva package from R ^74^.

Differential gene expression analysis was performed with R following the DEseq2 pipeline using the DEseq2 package ^75^. DEseq2 and edgeR – to retain only the expressed genes – algorithms were used. The packages are both available from Bioconductor (http://www.bioconductor.org) (accessed on October 2022). The threshold for significantly differentially expressed genes was set as an absolute log2 fold-change > 1. The volcano and correlation plots depicting the results were generated using the R package ggplot2 (Wickham, ggplot2: Elegant Graphics for Data Analysis, 2016). Gene-set enrichment analysis (GSEA) was performed using signatures from the literature, described in Supplemental Table 9, and expression obtained after DEseq2 normalization. GSEA parameters were set to 1,000 permutations per gene-set. Only gene sets with an NES > 1.7 and a FDR < 0.05 were considered.

### RNA quantification by RT-qPCR

RNA (3 µg) was reverse transcribed using M-MLV reverse transcriptase (Invitrogen) according to the manufacturer’s protocol. The newly synthesized cDNA was used as template for real-time quantitative PCR (qPCR) with the iTaq™ Universal SYBR Green Supermix. Technical triplicates were used for each sample and the quantified RNA normalized against *TBP* (human) or *Hprt* (mouse) as housekeeping transcripts (Tables S6 and S7).

### Immunofluorescence

Cells were seeded on coverslips and cultivated until 100% confluence was achieved. After 24 h of starvation in 0.5% FCS for murine melanoma cell lines and 1% FCS for human melanoma, the medium was complemented with 10 nM GRP and/or 1 µM RC for 1 h. Cells were fixed with 4% PFA for 15 min and blocked with 5% normal goat serum and 0.3% Triton™ X-100 in PBS. Cells were incubated overnight at 4°C with the anti-YAP D8H1X antibody (#14074, cell signaling, 1/100 dilution) in 1% BSA and 0.3% Triton™ X-100 followed by incubation for 1.5h at RT with goat anti-rabbit Alexa fluor 594 (#A-11012, Invitrogen, 1/500 dilution). Coverslips were mounted with Prolong® Gold containing 1.5 µg/mL DAPI. Images were acquired using an inverted SP8 Leica confocal microscope (Leica Microsystem). YAP1 localization was evaluated using ImageJ Software.

### Stimulation with estradiol

For hormone depletion, FCS was stripped using the dextran-coated charcoal method. Briefly, Norit activated charcoal (final concentration = 0.25%) and Dextran T-70 (final concentration = 0.0025%) in 0.25 M sucrose/1.5 mM MgCl2/10 mM HEPES, pH 7.4 were incubated overnight at 4°C. The volume equivalent to the volume of the FCS to strip was pipetted into a new 50 mL tube and centrifugated at 500 x g for 10 min to remove the supernatant. The FCS was incubated with the charcoal for 12 h at 4°C. The treated FCS was then filtered through a 0.22-µM pore filter to ensure sterility and mixed with the appropriate phenol-red free medium and PS. Cells were harvested using phenol red-free trypsin and starved for estrogen using phenol-red free/ 0% stripped FCS for four days. Cells were then stimulated with β-estradiol or ICI 182,780 for 72 h in phenol-red free/5% stripped FBS.

### Plasma stability assay

Each compound was diluted in mouse plasma to a final concentration of 1 μM and incubated at 37°C for 2 h. The reaction was stopped by the addition of 2.5 volumes ice-cold acetonitrile. Liquid chromatography-mass spectrometry was performed on the supernatant in multiple reaction-monitoring mode (LC-MS/MS). The percentage of the remaining test compound relative to that present at T0 was determined by monitoring the peak area.

### Metabolic stability assay

Each compound was diluted to a final concentration of 1 µM in 100 mM phosphate buffer (pH 7.4) containing 0.5 mg/mL mouse liver microsomes, 1 mM NADPH regenerating system, and 1 mM MgCl2 and incubated at 37°C for 1 h. At various time points (0, 2, 10, 20, 40, and 60 min), one volume of ice-cold acetonitrile was added and the supernatants analyzed by LCMS/MS. To obtain a stability curve, the percentage of remaining test compound at each time point was determined by monitoring the peak area. The half-life (T1/2) was estimated from the slope of the initial linear range of the logarithmic curve of the remaining compound (%) against time, assuming first-order kinetics.

### LC-MS/MS

Analyses were performed on a Shimadzu 8030 LCMS instrument. Chromatographic separations were carried out at 40°C using a 2.6-µm C18 Kinetex column (50 X 2.1 mm) purchased from Phenomenex. The mobile phase flow rate was set to 0.5 mL/min and the following program applied for the elution: 0 min, 5% B; 0-1.2 min, 5-95% B; 1.2-1.4 min, 95% B; 1.4-1.42 min, 95-5% B, and 1.42-2.8 min, 5% B (solvent A: 0.05% formic acid in water; solvent B: acetonitrile). The injection volume was 1 µL. The mass spectrometer was interfaced with the liquid chromatography system using an electrospray ion source. The nitrogen nebulizing gas flow was set to 1.5 L/min and the drying gas flow to 15 mL/min. The interface voltage was set to 4500 V. The temperature of the block heater was maintained at 400°C and the desolvation line at 250°C. Argon was used as the collision gas at 230 kPa. The transitions in positive mode were m/z 585.1 → 204.0, 221.0 for PD176252, and m/z 369.5 → 144.1, 110.1 for RC.

## QUANTIFICATION AND STATISTICAL ANALYSIS

Cell culture-based experiments were performed in at least biological triplicates and validated three times as technical triplicates. The significance of the effects was calculated using the Mann-Whitney test for the comparison of two groups or by ANOVA when comparing three or more conditions with an adjustment for multiple comparison using a Tukey test. Categorical data were compared using Fisher’s exact test when two groups were compared or, otherwise, the Chi^2^ test. The significance of the difference between Kaplan–Meier curves was calculated using the log-rank test. Data are represented as mean ± standard deviation unless otherwise indicated in the figure legend. All P values are reported as computed by Prism 6. P value < 0.05 were considered as significant. ns = not significant, ∗p < 0.05, ∗∗p < 0.01, ∗∗∗p < 0.001, ∗∗∗∗p < 0.0001.

### Data and code availability

Bulk RNA-seq data generated for this study have been deposited in the NCBI Gene expression Omnibus (GEO) (RRID:SCR_005012) and are publicly available at the data of publication (accessible in the SuperSeries GSE218588). Mouse tumor RNA-seq are available under accession number GSE218532. RNA-seq data for mouse and human cell lines are available under accession number GSE218586 and GSE218530 respectively. Any additional information required to reanalyze the data reported in this paper will be shared by the lead contact upon request.

### Supplemental information

#### Supplemental Figures

**Figure S1.**
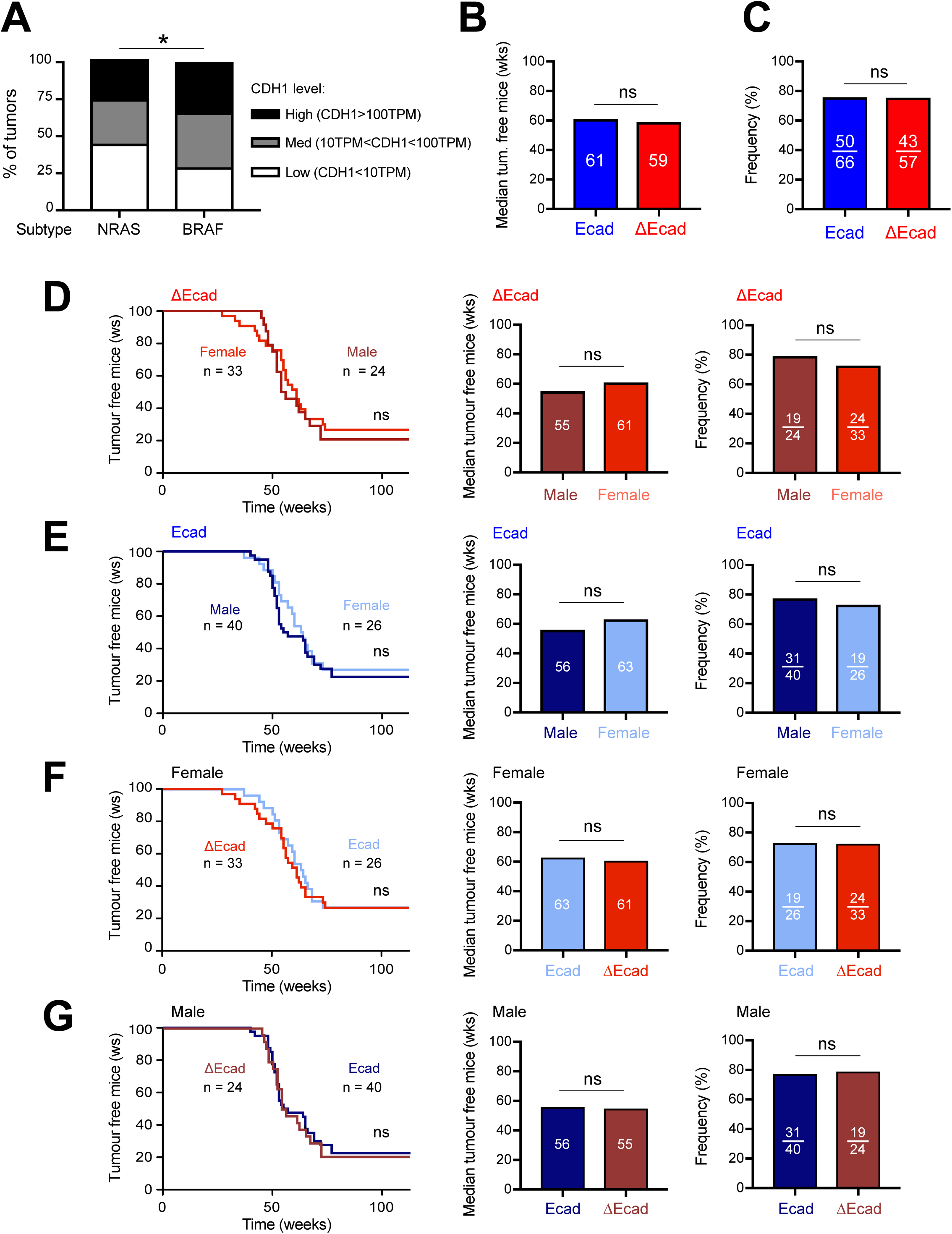
Effect of CDH1 expression associated with NRAS mutation in human and mouse melanoma. (**A**) Level of CDH1 expression according to the melanoma drivers NRAS or BRAF in the TCGA-SKCM database. Differences between the two populations were assessed using the Chi-square test. *p ≤ 0.05. (**B**) Median of tumor-free survival and (**C**) frequency of mice with primary melanoma according to the Ecad and ΔEcad genotype. (**D-G**) Kaplan-Meier, median, and frequency of tumor-free mice by genotype and sex. (**D**) Male versus female ΔEcad tumors, (**E**) male versus female Ecad tumors, (**F**) female ΔEcad versus female Ecad tumors, and (**G**) male ΔEcad versus male Ecad tumors. Kaplan-Meier curves were compared using the log-rank method. Statistical analysis of the frequency of tumor-free mice was performed using the two-sided chi-square test. ns: non-significant.

**Figure S2.**
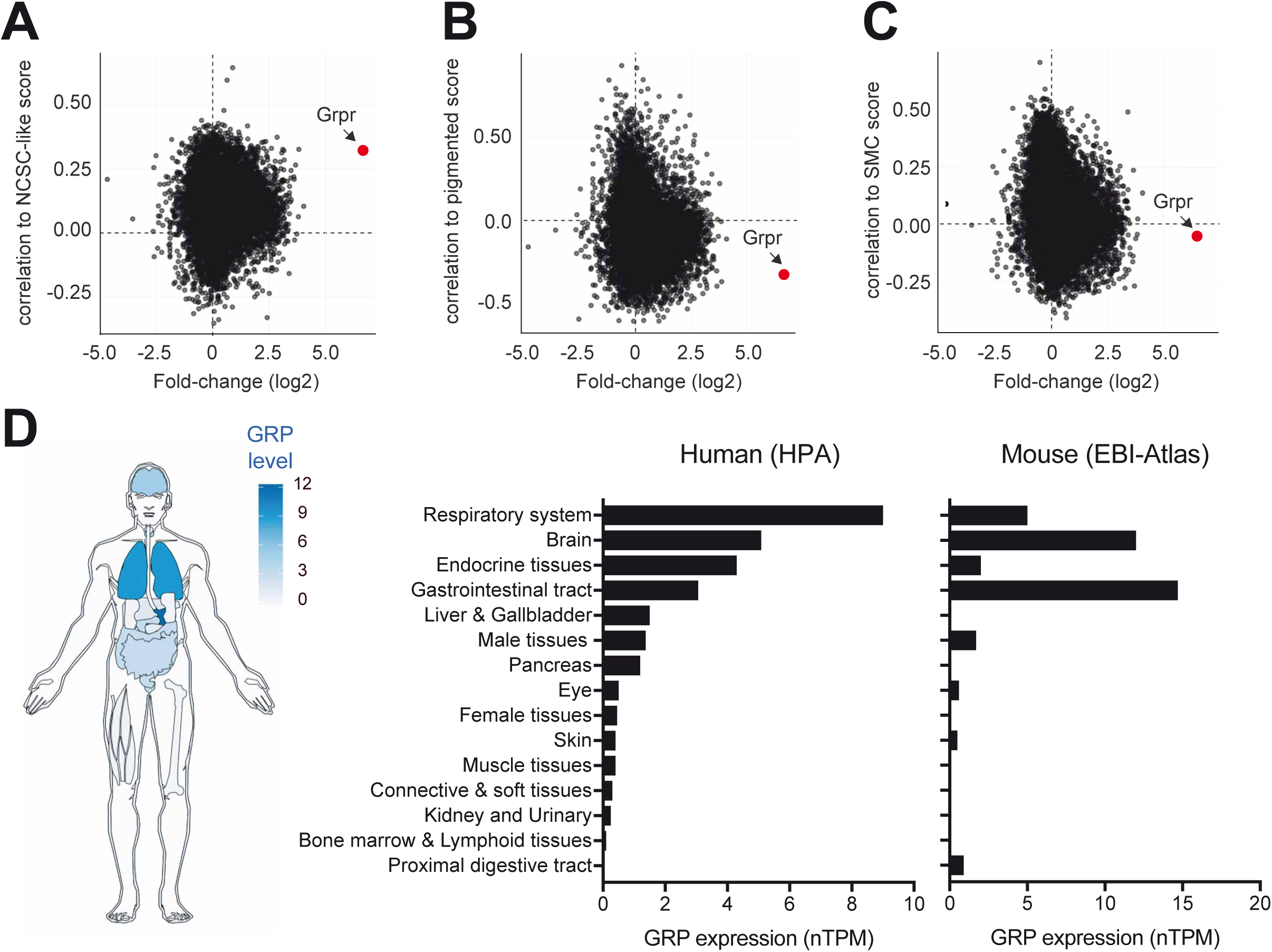
GRP, the endogenous GRPR agonist, is highly expressed in human and mouse lungs. (**A-C**) Plot of the differential gene expression (fold change, x-axis) with the Pearson correlation to phenotypic score (y-axis), (**A**) correlation to NCSC-like score, (**B**) correlation to pigmented score, (**C**) correlation to score SMC. The fold change in x-axis was calculated by the differential gene expression between female ΔEcad and female Ecad tumors. For each TCGA-SKCM tumor, each phenotypic score was calculated by averaging the expression of gene markers of the “NCSC”-like state, of the pigmented state and of the SMC state. The correlation of each genes expressed in the TCGA-SKCM tumors to each phenotypic score in y-axes was calculated using Pearson test. For A-C, Grpr is indicated in red. (**D**) GRP expression in humans and mice. Note that GRP is predominantly expressed in the human lung and highly produced in mouse lungs. The human data were extracted from the human protein atlas and the mouse data from the EBI expression atlas.

**Figure S3.**
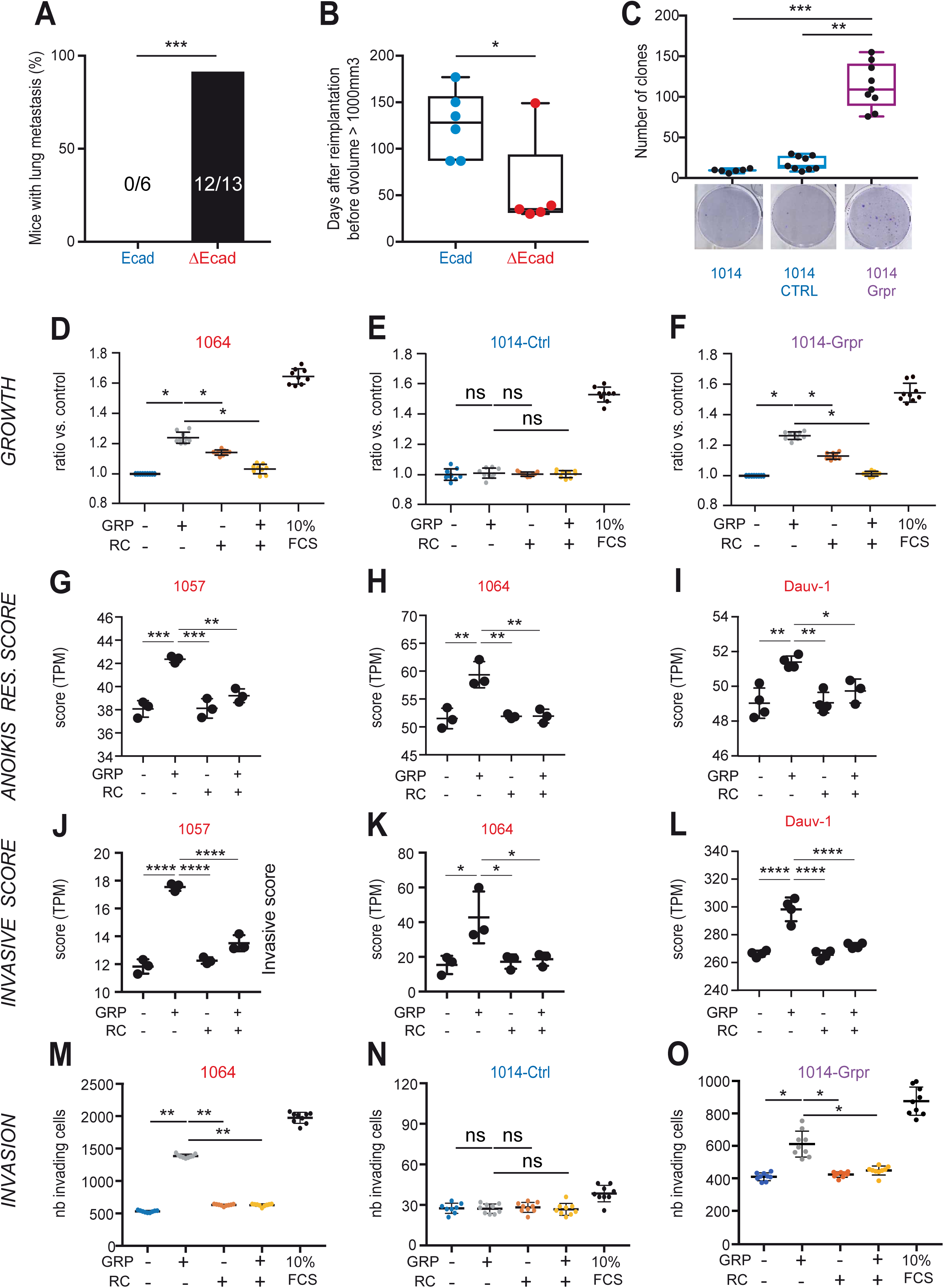
GRPR promotes the growth and invasiveness of melanoma cells. (**A**) Tail-vein injection of C57BL/6J mice with melanoma cells after dissociation from two independent Ecad and three independent ΔEcad tumors. Lung metastases were observed 50 days after injection in 12/13 injected mice with ΔEcad cells but were not observed with Ecad cells (0/6). Statistical analysis was performed using Fisher’s two-sided exact test. (**B**) Time after melanoma neck-graft reimplantation to reach a volume of 1 cm^3^ with female Ecad and ΔEcad tumor isografts. Statistical analysis was performed using a two-tailed Mann-Whitney test. (**C**) Clonogenic growth of the 1014 GRPR^neg^ melanoma cell line: left, parental cells; middle, control cell line; right, exogenous GRPR expression. Data are presented as the mean ± SD. (**D-F**) Effect of GRPR stimulation on the growth of the GRPR^pos^ mouse melanoma cell lines (**G**) 1064 and (**F**) 1014-Grpr and the GRPR^neg^ (**E**) 1014-Ctrl cell line. Cells were quantified for each condition. Data are presented as the mean ± SD. (**G-I**) Anoikis resistance score calculated from RNA-seq TPM-normalized data for the GRPR^pos^ murine (**G**) 1057 and (**H**) 1064 and human (**I**) Dauv-1 melanoma cell lines. Data are presented as the mean ± SD. (**J-L**) Invasive score calculated from RNA-seq TPM-normalized data for the GRPR^pos^ murine (**J**) 1057 and (**K**) 1064 and human (**L**) Dauv-1 melanoma cell lines. Data are presented as the mean ± SD. Number of GRPR^pos^ (**M**) 1064, (**O**) 1014-Grpr, and GRPR^neg^ (**N**) 1014-Ctrl murine melanoma invasive cells. (**D-O**) All assays were performed after overnight starvation. Cells were stimulated with vehicle (lane 1), 10 nM GRP (lane 2), 1 µM RC (lane 3), both (lane 4), or 10% FCS (lane 5, when present) for (**D-F**) 48 h, (**G-L**) 4 h, or (**M-O**) 24 h. (**M-O**) The invasion assay was performed in the presence of 200 µg/mL Matrigel®. Data are presented as the mean ± SD. Statistical analyses for D-O were performed by one-way ANOVA. Adjustments for multiple comparisons were performed using Tukey’s post-test.

**Figure S4.**
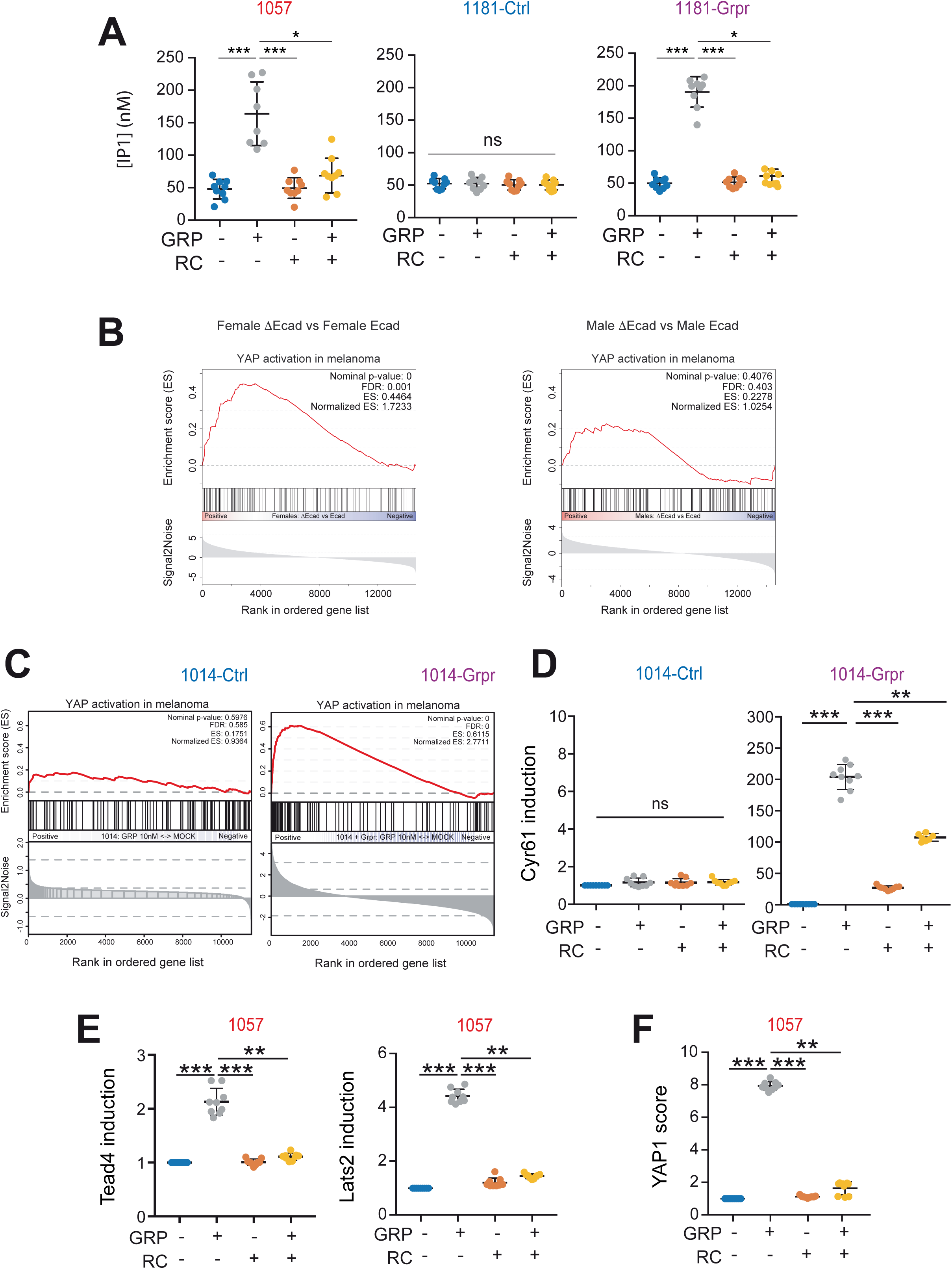
GRPR activation by GRP promotes the YAP1 transcriptional program. (**A**) IP1 concentration in the murine GRPR^pos^ 1057 (left), 1181-Grpr (right), and GRPR^neg^ 1181-Ctrl (middle) cell lines. (**B**) GSEA of the YAP1 activation signature in murine primary melanoma tumors comparing ΔEcad female tumors to Ecad female tumors (left) and ΔEcad male tumors to Ecad male tumors (right). mRNA expression was analyzed by RNA-seq and the data normalized using DEseq2 prior to conducting the GSEA. (**C**) GSEA analysis of the YAP1 activation signature in murine melanoma cell lines expressing Grpr (1014-Grpr) or not (1014-Ctrl) 4 h after stimulation with 10 nM GRP. mRNA expression was analyzed by RNA-seq and the data normalized using DEseq2 prior to conducting the GSEA. (**D**) Induction of the Yap1 target Cyr61 in the 1014-Ctrl (left) and 1014-Grpr (right) murine melanoma cell lines. (**E**) Induction of Yap1 targets in the 1057 GRPR^pos^ melanoma cell line: Tead4 (left) and Lats2 (right). (**F**) Quantification of the Yap1 score in the GRPR^pos^ 1057 melanoma cell line. (**A,D-F**) cells treated with vehicle (lane 1), 10 nM GRP (lane 2), 1 µM RC (lane 3), or both (lane4). Data are presented as the mean ± SD. Statistical significance for (**A,D-F**) was calculated by ANOVA followed by an adjustment for multiple comparisons using Tukey’s post-test.

**Figure S5.**
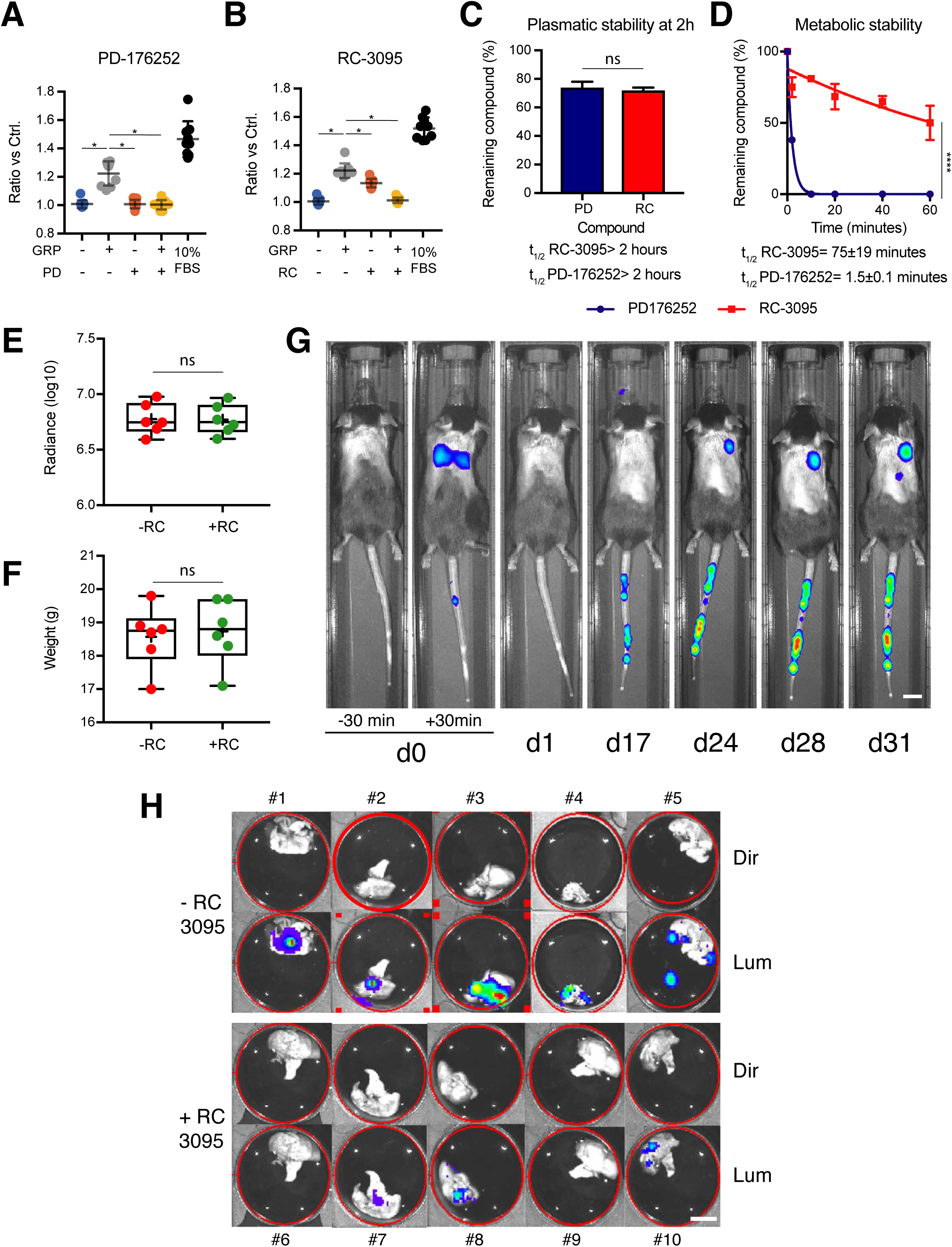
RC inhibits the growth and number of lung melanoma metastases. (**A-B**) The GRPR inhibitors RC-3095 (RC) and PD-176252 (PD) have the same blocking effect on GRP-induced growth. The GRPR^pos^ 1057 melanoma cell line was starved overnight and then treated in low-serum medium containing vehicle (DMSO 0.1%, lane 1), 10 nM GRP (lane 2), 1 µM GRPR inhibitor (lane 3), 1 µM GRPR inhibitor + 10 nM GRP (lane 4), or 10% FBS (lane 5). The GRPR inhibitor was (**A**) PD-176152 or (**B**) RC-3095. Growth was assessed by MTT 48 h after treatment. Significance was measured by ANOVA, adjusting for multiple comparisons using Tukey’s post-test. (**C**) Ex vivo stability in murine plasma and (**D**) metabolic stability after incubation with mouse liver microsom7es of RC (red) and PD (blue). The decay curves for panels **C** and **D** were modeled from the data based on an exponential decay curve equation. The t1/2 (time when the concentration is reduced by half) was calculated from the obtained equations. (**E**) Mean weight and (**F**) thorax radiance 30 min after cell injection for the RC-treated and vehicle-treated groups after randomization. (**G**) Representative images of C57BL/6 mice intravenously injected with 1057-Luc melanoma cells analyzed using the IVIS Spectrum Imaging System at various time points (d = day). Note that at day 31, a signal was detected in other organs. After dissection, metastases were observed in the liver, adrenal glands, and ovary (see signal appearing at day 31 outside of the lungs). Scale bar= 1 cm (**H**) Detection of luminescence by IVIS imaging of the lungs after euthanasia and dissection of RC-treated (n#6 to #10) or untreated (n#1 to #5) mice. Scale bar= 1 cm Lum: luminescence, Dir: direct.

### Supplemental Tables

#### Table titles and legends

Table S1. Differential expression analysis results of female ΔEcad tumors versus female Ecad tumors. Eight female ΔEcad tumor and eight female Ecad tumor transcriptomes were compared by RNA-seq using DEseq2.

Table S2. Differential expression analysis results of male ΔEcad tumors versus male Ecad tumors. Eight male ΔEcad tumor and eight male Ecad tumor transcriptomes were compared by RNA-seq using DEseq2.

Table S3. Differential expression analysis results of female Ecad tumors versus male Ecad tumors. Eight female Ecad tumor and eight male Ecad tumor transcriptomes were compared by RNA-seq using DEseq2.

Table S4. Differential expression analysis results of female ΔEcad tumors versus male ΔEcad tumors. Eight female ΔEcad tumor and eight male ΔEcad tumor transcriptomes were compared by RNA-seq using DEseq2.

Table S5. Cell line characteristics. Origin and detailed genotypes of each cell line and associated expression level of key genes assessed by RNA-seq.

Table S6. Primers used Table S7. PCR conditions Table S8. siRNA

Table S9. GSEA signatures

